# Sphingosine-1-phosphate regulates Plasmodium histone deacetylase activity and exhibits epigenetic control over cell death and differentiation

**DOI:** 10.1101/2022.06.09.495418

**Authors:** Raj Kumar Sah, Sakshi Anand, Geeta Kumari, Monika Saini, Rahul Singh Hada, Evanka Madan, Aashima Gupta, Shailja Singh

**Affiliations:** Special Centre for Molecular Medicine, Jawaharlal Nehru University, New Delhi: 110067, India; Department of Life Sciences, School of Natural Sciences, Shiv Nadar University, Greater Noida-201314, India

**Keywords:** Malaria, Plasmodium, Histone deacetylase 1, Sphingosine 1 Phosphate, Multistage

## Abstract

Histone deacetylases (HDACs) play a key role in cellular processes by the regulation of gene transcription. This study contributes a novel insight how *Plasmodium falciparum* HDAC (*Pf*HDAC-1) is regulated by S1P produced by host erythrocyte SphK-1. The binding of S1P with endogenous nuclear extract *Pf*HDAC-1 and recombinant *Pf*HDAC-1 blocks their activity. A significant modulation in transcriptional regulation of *P. falciparum* HDAC regulated genes resulted upon inhibition of S1P production through blocking of hSphK-1 by clinical SphK-1 inhibitor PF-543. PF-543 led to profound decrease in S1P in the parasite’s nuclear fraction. The significant modulation of *Pf*HDAC-1 regulated specific candidate genes related to gametocytogenesis, virulence and proliferation was observed in parasite treated with SphK-1 inhibitor, suggesting S1P targets *Pf*HDAC-1 and participates in epigenetic regulation of these key cellular processes. The epigenetic modulation of parasite cell growth and differentiation by host provides a novel approach for the developmenthost-targeted therapeutics.

## Introduction

In 2020, one year after the COVID-19 epidemic and service delays, the anticipated number of malaria cases soared to 241 million ^1^. In Southeast Asia and Africa, *P. falciparum* has developed resistance to all antimalarial medicines, including artemisinin-based combination regimens^23^. In order to combat malaria, drug development is essential. First-line antimalarial treatments are mainly targeted at asexual blood-stage parasites^6, 45^. To avoid disease outbreaks and protect susceptible populations from asexual drug resistance, it is necessary to prevent the liver- and gametocyte-stage parasites^7, 8^.

Plasmodium falciparum has a lot of genetic and phenotypic latitude, which helps it spread malaria. Despite its early evolutionary position, the parasite has made large genome-use investments^9^. During its 48-hour intraerythrocytic development cycle, the parasite coordinates the expression of essential genes, undergoing periods of enhanced metabolic activity and replication before infecting new cells^10–12^. In addition, it has been demonstrated that various signaling molecules, such as Ca2+ and cAMP, regulate the release of essential components involved in the life cycle of the human malaria parasite. When crucial molecules are released, parasites enter host erythrocytes, reproduce, exit, and re-enter red blood cells to continue their life cycle^13–15^. The *P.falciparum* epigenome is linked to the regulation of several important parasite processes, including the control of parasite proliferation during asexual development and sexual differentiation^16, 17^. Uniqueness of the epigenomehas prompted research into new chemotypes that target epigenetic modulators^16^. Since the discovery of the first epigenetic enzymes controlling histone acetylation for transcriptional regulation of genes, numerous inhibitors of Since the discovery of the first epigenetic enzymes controlling histone acetylation for gene transcription, several inhibitors of epigenetic enzymes, particularly histone deacetylases (HDACs), have been created for clinical use against many diseases, including cancer^18, 19^. Histone changes such as acetylation, deacetylation, and methylation regulate essential malaria parasite pathways^20^. According to prior studies, three of the five *P. falciparum* HDAC (*Pf*HDAC) genes in the parasite genome are essential for the parasite’s blood stage ^21, 22^, and HDAC classifications are based on sequence matching to yeast prototypes. Classes I, II, and IV are zinc-dependent yeast deacetylases^20, 23–25^. *P. falciparum* has Class I and Class III HDAC homologues, *Pf*HDAC1 and *Pf*Sir2^26, 27^. *Pf*HDAC1 transcripts were found in asexual and exo-erythrocytic stages. *Pf*Sir2 connects with telomeric clusters, causing heterochromatin. *Pf*Sir2 binding and deacetylation affects telomeric var genes’ surface antigen expression^20, 28, 29^. Numerous studies have identified potent anti-plasmodial HDAC inhibitors, including dual-/multistage-targeting compounds, implying that *Pf*HDACs could be targeted in drug development^7^. In recent study, researcher have characterized the class I histone deacetylase *Pf*HDAC-1 and demonstrated that phosphorylation of *Pf*HDAC-1 is required for its catalytic activity^30^. *Pf*HDAC-1 binds to and regulates parasite genes responsible for housekeeping and stress-responsive functions. Some drugs have also been shown to have direct effects on histone acetylation, deacetylation, and transcriptional alterations. Direct evidence that *Pf*HDACs are the target of these drugs is absent^7^.

As parasites and cancer cells rely more on lactate fermentation as an energy source, they may share several traits, including the ability to evade the immune system^31–33^. The epigenome, which includes DNA methylation and histone changes, has been extensively studied to uncover novel targets in drug discovery programmes ^34–36^. Plasmodium falciparum and vivax have a continuous gene expression cascade during intraerythrocytic development (IDC). Most genes are activated only once every cycle, when their products are needed^20, 37, 38^. Histone acetyltransferases (also known as HATs) and histone deacetylases coordinate to maintain the delicate, dynamic balance of nucleosomal histone acetylation levels^39^.

Compared to PTM, there isn’t much evidence that lipids regulate cell epigenetics (PTM). Nuclear lipid metabolism has changed how we think about phospholipids as signalling molecules^40^. Inositol lipids are the best-characterized intranuclear lipids are important for pre-mRNA splicing, mRNA export, transcriptional regulation, and chromatin remodelling^41^. Sphingomyelin is well-known in nuclear matrix. Recent evidence suggests that sphingolipids are metabolised in the nucleus^41–43^. HDAC1 and HDAC2 are two human histone deacetylases recognised for their enzymatic activity, which has been shown to be regulated by S1P, which prevents the removal of acetyl groups from histone residues within the histone tail^41^. It has been shown that the plasmodium sphingolipid metabolism pathway involves several enzymes (e.g., glucosylceramide synthase, two sphingomyelinases)^44–46^. An enriched host cell lipid pool may act as a reservoir for parasite-mediated salvage. Several studies have suggested that *P. falciparum* may import certain types of lipid, including ceramides, sphingolipids, and Lyso PCs^47, 48^. In our previous study, it has been found that plasmodium does not contain any SphKs enzyme; as a result, it uptake S1P from the host pool ^49, 50^. Learning more about this very specialised form of transcriptional control will be intriguing. According to growing findings from eukaryotes, histone alterations affect chromatin structure and gene expression^51, 52^. In light of this, we hypothesized that *P. falciparum* scavenges sphingolipid metabolites for intracellular proliferation and growth via epigenetic reprogramming mediated by S1P.

Consistent with this theory, we identified that S1P interacts with PfHDAC-1, as determined by microscale thermophoresis (MST), dot blot and immunoblot analysis, and regulates the activity through histone acetylation state. *P. falciparum* is examined for histone acetylation and hyperacetylation assays and HDAC activity assays in order to comprehend S1P-mediated HDAC activity. PF-543, a particular inhibitor, was employed to better comprehend the function of S1P by reducing its level in infected erythrocyte. Next, we examined the transcriptional impact of decreased S1P by measuring the expression of *Pf*HDAC-1-regulated genes. Further, the production of alpha tubulin is postulated as a potential transcriptional marker of HDAC inhibition in malaria parasites, which is consistent with the findings of the study headed by ^30^. The transcriptome analysis of *Pf*HDAC-regulated specific candidate genes related to gametocytogenesis, virulence, and proliferation was reduced in parasites treated with SphK-1 inhibitor, indicating that S1P targets *Pf*HDACs and participates in epigenetic regulation of gene expression of *P. falciparum* in relation to progression, sexual commitments, and virulence. To show the effect of S1P by PF-543 inhibition in *in vivo P.berghie* infected mice modal was used to decipher the role of S1P in growth and development of sporogonic cycle in *P.berghie* infected mice, This work which was undertaken for the first time on *P.falciparum*, demonstrates how parasites acquire host pool lipid to modify epigenome. This research provides new insights into how *Pf*HDAC is regulated and how it contributes to S1P-mediated epigenetic regulation, both of which provide tremendous insight into the development of effective drugs for limiting the development of the malaria parasite.

## Results

### Recombinant PfHDAC1 interacts with S1P

Despite the fact that PfHDAC1 has been reported as a target of several very effective inhibitors, regulation, and biological characterization of PfHDAC1 function in *P. falciparum* is not yet established but recent publication from ^30^ karmodiya et al. shows the regulation of HDAC-1 activity via phosphorylation and how HDAC regulate many genes related to virulence, gametocyte etc. S1P preferentially binds to and inhibits histone deacetylases HDAC1 and HDAC2, restricting the removal of acetyl groups from lysine residues in mammalian histone tails^41^. We cloned, expressed, and purified histidine tagged PfHDAC1 to investigate its enzymatic activity (**Fig. 1a and 1b, supplementary Fig.1**). We sought to determine whether S1P interacted with PfHDAC-1. Initially, we generated an antibody against *Pf*HDAC1 the full-length protein and evaluated its specificity using immunoblotting with recombinant protein and parasite lysate (**Fig. 1c**). The immunoblotting results for *Pf*HDAC-1 exhibit a distinct band. In addition, a microscale thermophoresis (MST) experiment was conducted to confirm the binding of S1P to HDAC-1. We examined the thermophoresis change after titrating various concentrations of S1P to HDAC-1. MST confirmed the in silico binding of S1P and positive control SAHA to *Pf*HDAC-1 (**Fig. 1 d,e**). For both ligands, concentration-dependent fluorescence signals were measured, indicating an interaction between both ligands and the fluorescently labeled *Pf*HDAC-1 protein. The calculated dissociation constant for S1P (Kd = 177nM ± 24nM) indicated a stronger binding of S1P to PfHDAC-1 than in the case of SAHA (Kd = 2.09μM± 0.45 μM). For both ligands, concentration-dependent fluorescence signals were measured, indicating an interaction between both ligands and the fluorescently labeled hHDAC-1 protein. The calculated dissociation constant for S1P (Kd = 1.61μM ± 0.8μM) indicated a stronger binding of S1P to PfHDAC-1 than in the case of SAHA (Kd = 15.06μM ± 0.75μM). Altogether these results indicate the binding of S1P with PfHDAC-1 which could be endogenous ligand for HDAC-1.

**Figure 1:**
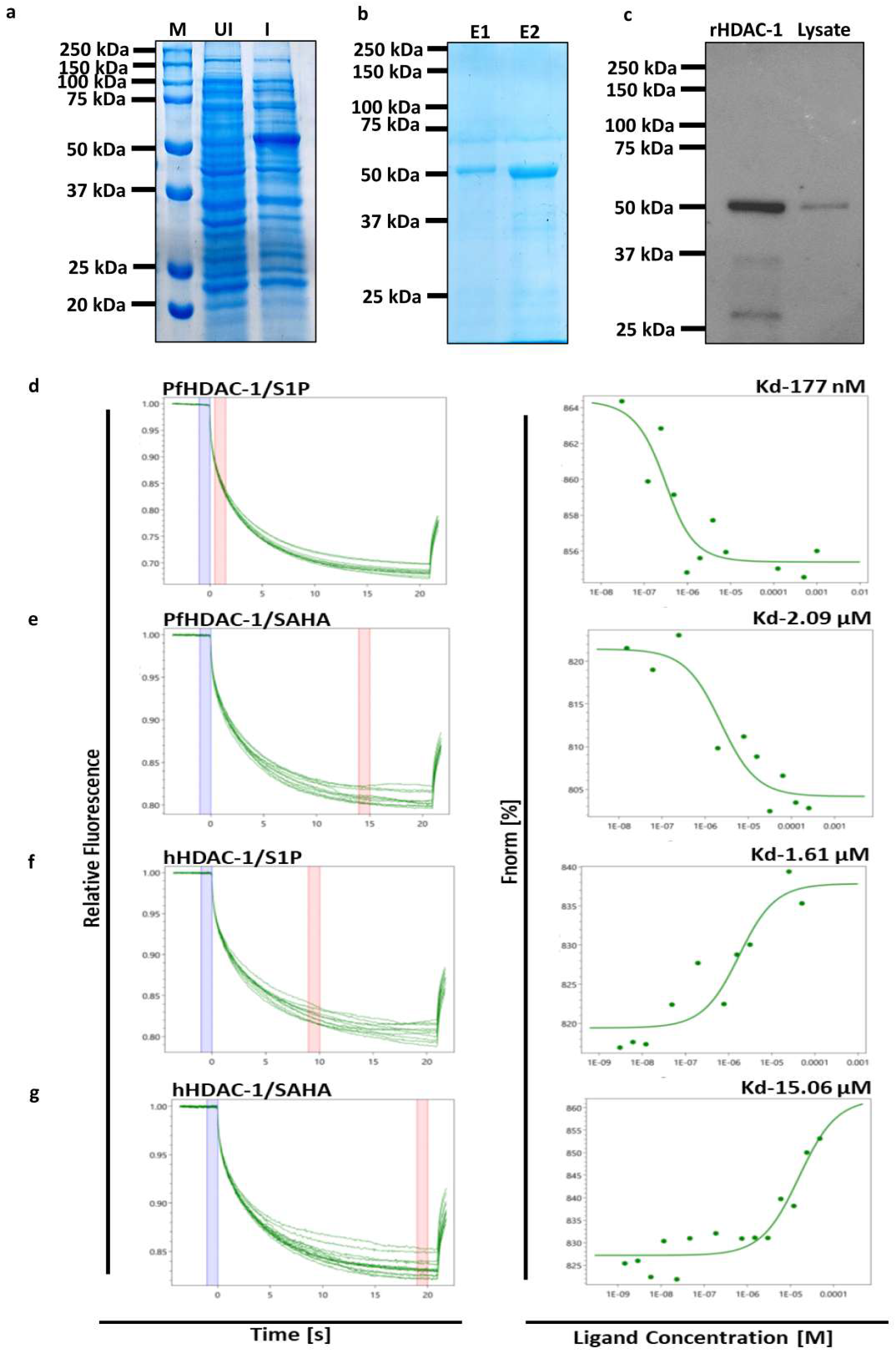
S1P interacts with recombinant *Pf*HDAC1. **a-b)**. SDS-PAGE gel showing induction and purification of recombinant PfHDAC-1 expressed in *E.coli* BL21 with an expected size of 55kDa. **c)** Immunoblot detection of purified recombinant PfHDAC1 and endogenous PfHDAC1 across asexual blood stages using mice anti-rPfHDAC1 polyclonal antibody **d-g).** Microscale thermophoresis-based protein-protein interaction assay between **d)** S1P-PfHDAC1, **e)** SAHA-*Pf*HDAC1, **f)** S1P-hHDAC1 and **g)** SAHA-hHDAC1. Binding of S1P to fluorescently labeled *Pf*HDAC1 and hHDAC1 measured by MST resulted in a *K_d_* value of 177nM and 1.61μM respectively. Whereas, for SAHA-PfHDAC1 and SAHA-hHDAC1 complex formation, a *K_d_* value of 2.09μM and 15.06μM was obtained.

### Host S1P interacts with HDAC-1 and reconfigures its activity, as determined by in silico and deacetylase activity

*Pf*HDAC-1 protein, containing multiple beta sheets and helices, consisting of 449 residues. To evaluate the physical basis of protein-ligand interactions, we performed *in-silico* docking studies of PfHDAC-1proteinwith SAHA and S1P compounds. The best conformations of the docked compounds were selected based on their lowest free binding energy to the binding domain. Analysis of this pocket unravelled one hydrophobic interaction with TYR301 (substrate binding site), five residues HIS139 (Active site; proton acceptor), GLY147(substrate binding site), ASP174, HIS176 (metal binding site) and TYR301 with hydrogen bond interactions using minimum binding energy of -6.63kcal/mol (**Fig. 2b), Supplementary Table 4)**. Likewise, S1P interacts with PfHDAC1 with seven residues involved in hydrophobic interaction and four residues ALA22 (2 bonds), GLU94 (2 bonds) involved in H-bond interaction using minimum binding energy of -5.74kcal/mol (**Fig. 2a, supplementary Table 2.**). Further, we performed the docking of S1P with hHDAC1 and hHDAC2 proteins. We found that S1P interacts with hHDAC1 with three residues involved in hydrophobic interactions and five residues involved in H-bond interaction with minimum binding energy of -7.71kcal/mol for hHDAC1 (**Fig. 2c Supplementary Table 3**). Similarly, we found that S1P interacts with hHDAC2 with seven residues involved in hydrophobic interactions and seven residues **(Supplementary Table 4)** involved in H-bond interaction with minimum binding energy of -8.65 kcal/mol for hHDAC2 (**Fig. 2d**). Analysis of these results suggested that S1P forms stable structure with HDAC binding pocket. We compared hHDAC1 and hHDAC2 to Plasmodium falciparum isoforms. hHDAC1 and hHDAC2 revealed 60% similarity with PfHDAC1 and 25%-35% identity with PfHDAC2 as mentioned in **supplementary Table 5, and supplementary fig. 4,5.** Recently, in our previous study we have shown that parasite uptake host derived S1P from host. The binding interaction between HDAC-1 and S1P was further confirmed by performing lipid dot blot assay. For this assay, 500ng, 250ng, 125ng and 87.5ng of S1P were spotted on the nitrocellulose membrane (**Fig. 2e**). The dark spots of varying size that formed against each dot of S1P when incubated with rPfHDAC-1 and rPfH4, confirmed the binding between HDAC1 and S1P while no such spot pattern was observed when r*Pf*H4 was incubated with S1P.

**Figure 2:**
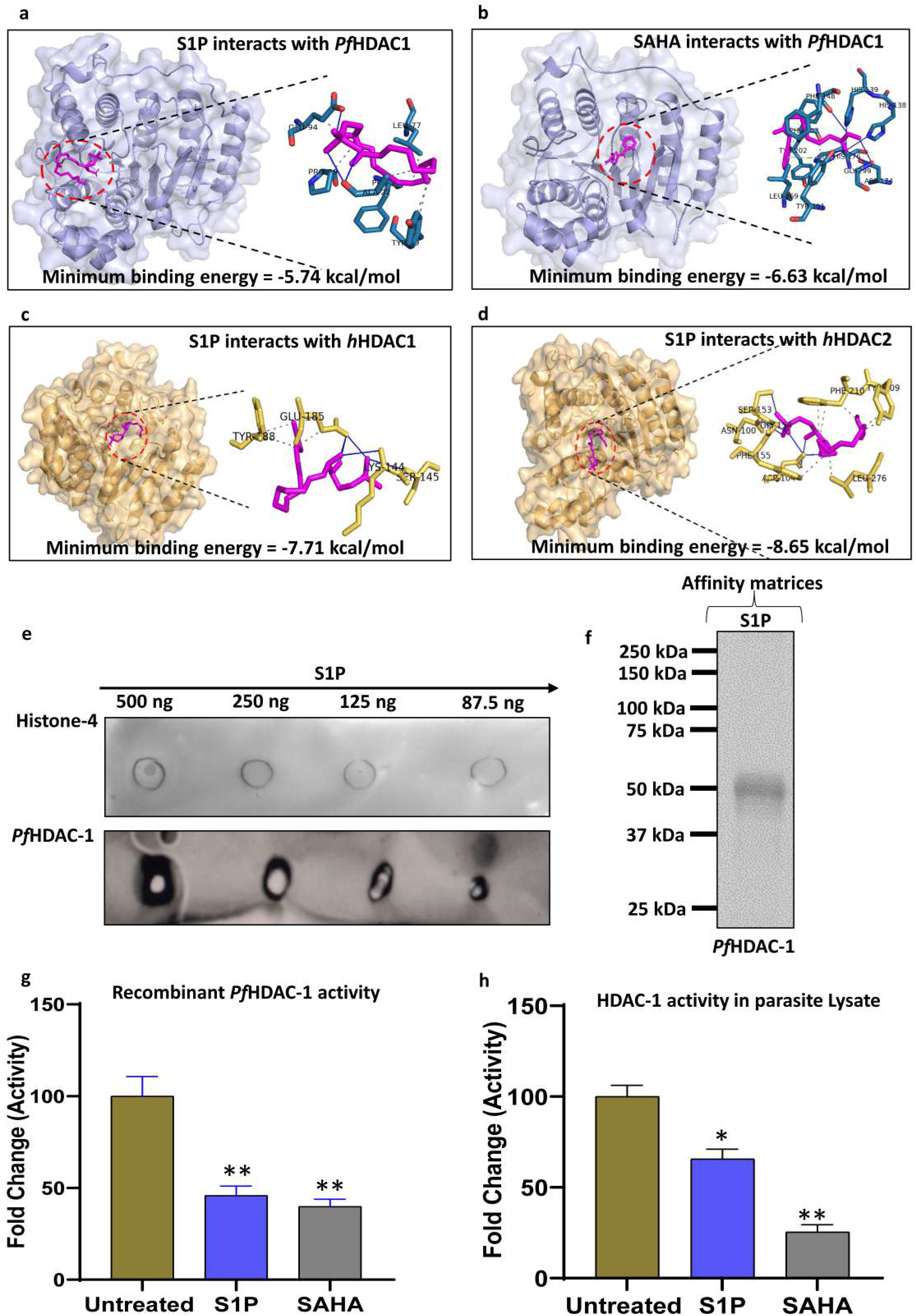
Validation of HDAC-1 binding with S1P and activity. **a-d)** Docking based protein-ligand interaction studies were done to show the binding interaction of S1P with PfHDAC1 and S1P with hHDAC1/hHDAC2. The known binder of HDAC1, SAHA was also used as a positive control for the study. **e)** Lipid dot blot with the indicated amounts of S1P spotted onto Nitrocellulose Membrane filter, showed binding of rPfHDAC-1with S1P as detected by probing with anti-His antibody. No significant binding was observed with the negative control rPfH4. Quantification of the level of rPfHDAC-1 and rPfH4 binding to different concentrations of lipid was determined from the integrated densities of spots on immunoblots. Values are given as arbitrary units (mean ± SEM; n = 3 independent experiments. **f)** Pull-down of rPfHDAC by S1P agarose beads. Elute obtained after washing of S1P beads incubated with protein was subjected to SDS-PAGE and the detection was confirmed using immunoblot assay. **g-h)** Fluorimetric activity assay of HDAC. Nuclear lysates of *Pf*3D7 parasite and the rPfHDAC protein were incubated with S1P (20 μM), positive control SAHA (2 μM) and a no enzyme control. The fluorescence was read by setting the excitation and emission wavelength of 340-360 nm and 440-465 nm, respectively. Graph depicting the fold change in the activity of HDAC was plotted. Bar graphs represent the relative fluorescence units (RFU) measured after 30 min. (n= 3 independent experiments, \**p* ≤ 0.05).

We pulled down S1P-binding proteins using a quick and sensitive approach. A distinct protein band with an apparent molecular mass of ∼52 kDa was evident in the pulldown of nuclear proteins with S1P-conjugated beads that was not present in the absence of nuclear extract or when control unconjugated agarose beads were used (**Fig.2f**). Western blot analysis with a specific PfHDAC-1 antibody confirmed the identity of this S1P-binding protein (**Fig.2f**). Pull down and dot blot analysis confirmed the interaction of S1P with PfHDAC-1 followed by in silico analysis. Further to check whether S1P interaction with PfHDAC-1 has any impact on its activity we validate by quantify the activity of *Pf* HDAC1, a fluorescence based HDAC activity assay was performed. Since it was evident from our data that S1P binds to HDAC1, we speculated that this binding interaction will possibly result in diminution of HDAC1 activity. Certainly, a 50% reduction in HDAC1 activity upon treatment with S1P was observed as indicated by the difference in fluorescence intensities of control and treated samples. The known inhibitor of HDAC1, SAHA was taken as the positive control. Similar results were observed with both recombinant protein and parasite nuclear lysate (**Fig. 2g,h**). Altogether these observations indicate the interaction of S1P with PfHDAC-1 and S1P modulate the activity of PfHDAC-1.

### The inhibition of host SphK-1 activity by the specific inhibitor PF-543 reduced S1P levels in parasite nuclei

Previous reports suggested that protozoa and fungal pathogens utilize sphingolipid metabolites during host pathogen interactions17,36,37. As a result, we used the SphK-1 specific inhibitor PF-543 to evaluate the level of S1P in parasite-infected erythrocytes. We made protein his tagged SphK-1 and check its activity and binding affinity with PF-543 to evaluate the inhibitor in *in vitro*. Docking study and MST analysis of SphK-1 and PF-543 shows good binding affinity and activity inhibition in PF-543 treatment **(Fig.3a-e, Supplementary figure 6)**. Further PF-543 a specific inhibitor of SphK-1 was used to downregulate the activity and checked at different time point in parasite nuclei. The bar graph represents the level of NBD-S1P in nuclei after PF-543 treatment at 4 and 12h done by fluorimetry method **(Fig.3f)**. Then we validated the intracellular localisation of NBD-S1P in infected erythrocytes and saponin lysed parasite by using fluorescence microscopy. Fluorescence micrograph and line graph analysis clearly suggest the uptake and localisation of NBD-S1P in nuclei as data indicate localisation with DAPI in infected erythrocytes and host free parasite after saponin lysis (**Fig. 4a, b**).

**Figure 3:**
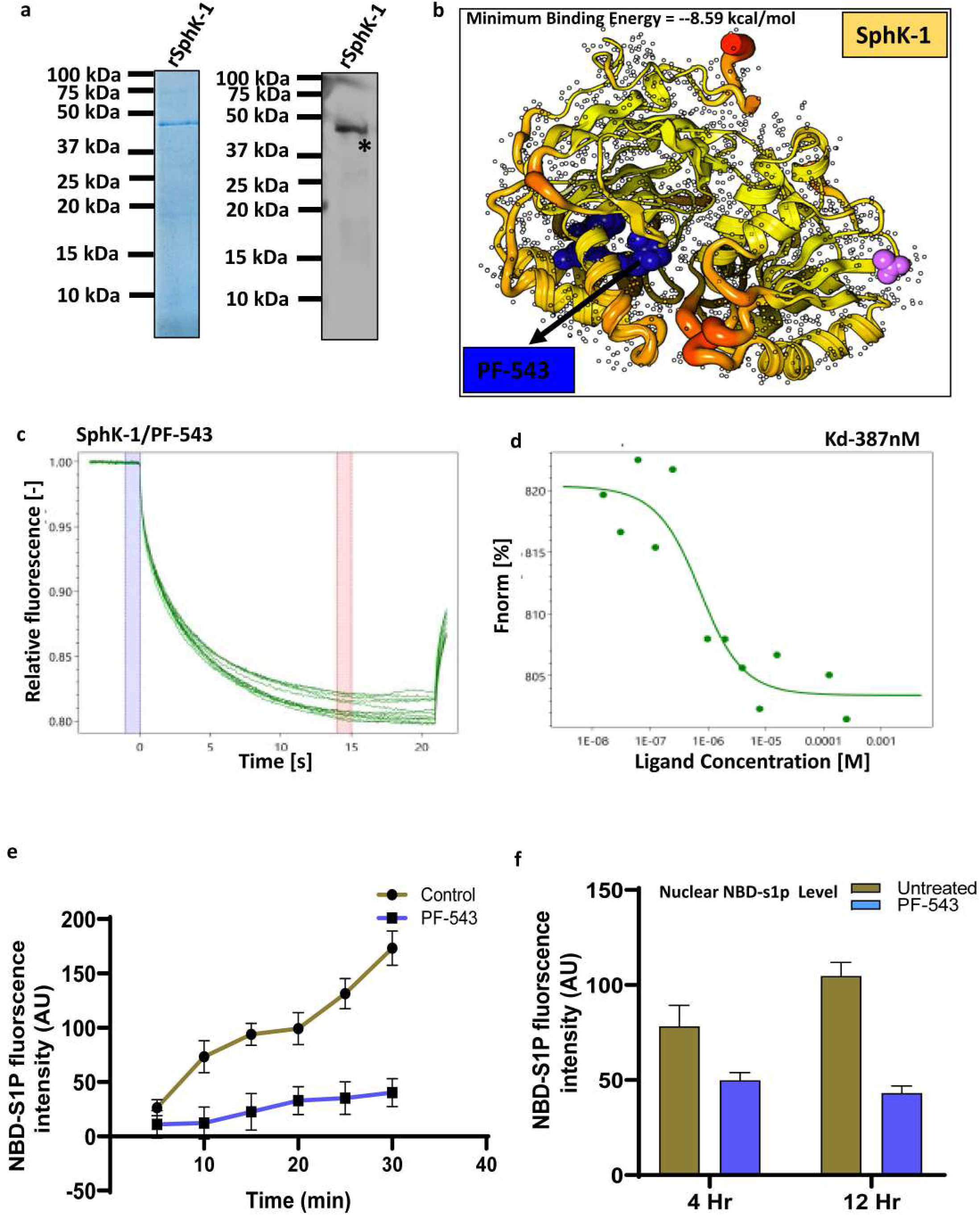
**a)** SDS-PAGE gel showing purification of recombinant hSphk-1 expressed in *E.coli* BL21 with an expected size of 45 kDa and its Immunoblot detection using mice anti-hSphk1 polyclonal antibody **b)** Molecular docking analysis to show the binding interaction between hSphk1 and PF-543**. c-d)** Microscale thermophoresis based protein-protein interaction assay between hSphk1 and PF-543. The binding resulted in a *K_d_* value of 389 nm indicating the high binding specificity of PF-543 for hSphk1. **e**-**f)** Intensity plot and Bar graph showing fluorimetric analysis of NBD-S1P level in parasite nuclei upon treatment with PF-543 at different time points and untreated control. (n=3 independent experiments, \**p* ≤ 0.05)

**Figure 4:**
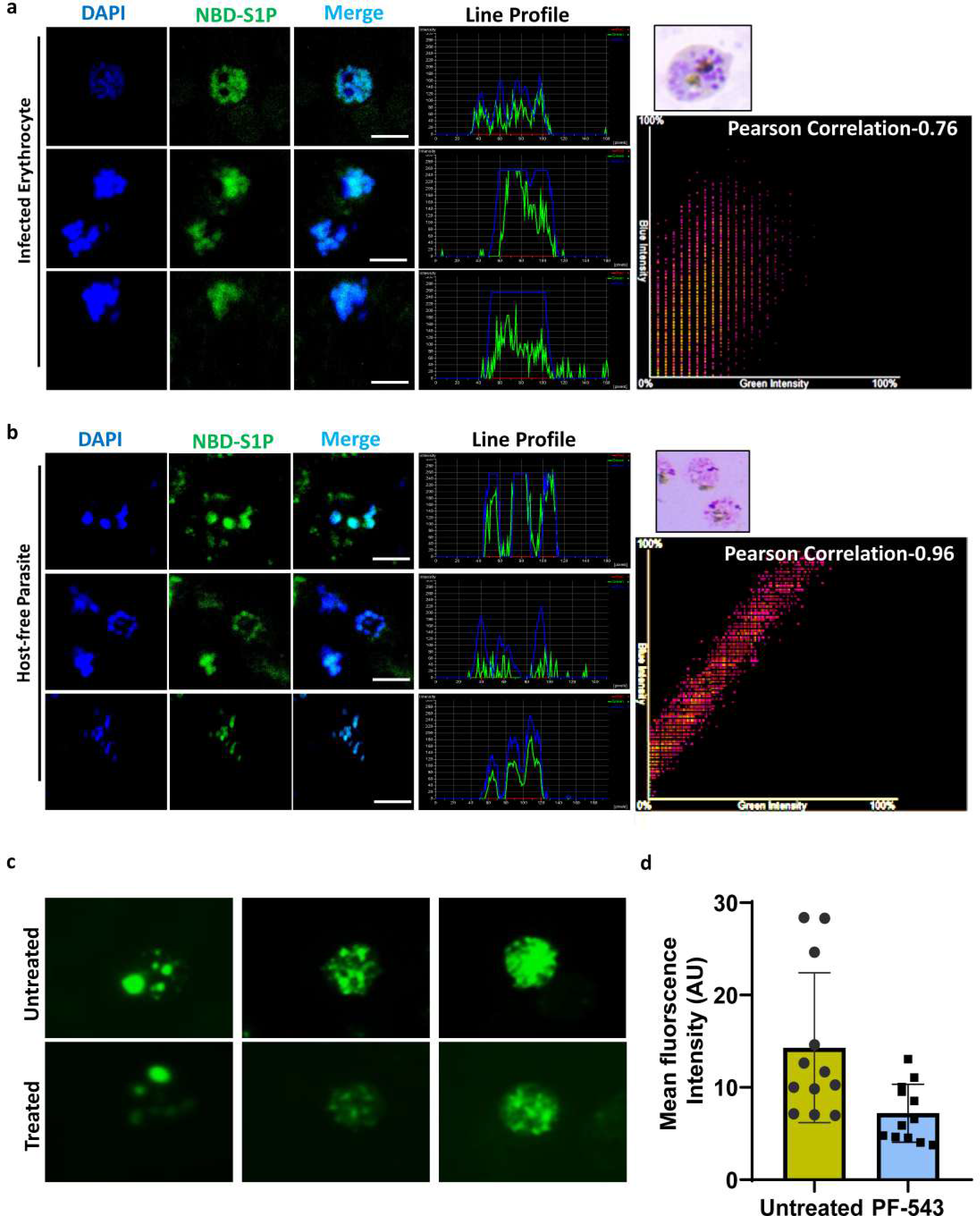
Corroboration of host S1P uptake by parasite. **a-b)** Immunofluorescence assay of infected erythrocytes and host-free parasites exhibiting the localization of NBD-S1P in parasite nuclei. The uptake of NBD-Sph by infected erythrocytes and its conversion into NBD-S1P followed by incorporation of the same in parasite nuclei is depicted by the co-localization of NBD-S1P with nucleus stained with DAPI. Scale bar =5μM **c)** Difference in fluorescence intensities of NBD-Sph in untreated control versus PF-543 treated parasites. **d)** Bar graph manifests significant reduction in NBD-S1P formation due to PF-543 mediated inhibition of SphK1. (n=3 independent experiments, \**p* ≤ 0.05)

The results showed that parasites isolated from infected erythrocytes with 30 min incubation with NBD-Sph had significant NBD-S1P uptake, which was reduced to 50% following treatment with PF-543, suggesting parasite co-dependency to host S1P pool. Fluorescence and quantification of NBD-S1P intensity plotted as bar graph (**Fig.4 c, d**). This finding revealed that s1p localised to nuclei as observed by fluorometric and microscopy analysis. PF-543 inhibited S1P synthesis in infected erythrocytes, as seen by decreased fluorometric intensities of fluorescent tagged NBD-S1P.

### In cell analysis of S1P interaction with HDAC-1 and Inhibiting SphK-1 in the host regulates S1P-mediated histone acetylation

Immunoblotting analysis was done to examine the effect of the S1P and HDAC inhibitor on H4 acetylation in *P. falciparum* 3D7 parasites **(Fig.5a)** The acetylation signals obtained after probing with anti-(tetra)-acetyl histone H4 antibodies were significantly increased for the HDAC inhibitor (SAHA) and decreased for the PF-543 treatment relative to the untreated control, histone-4 used as a loading control **(Fig.5a)**. Histone acetylation antibodies (anti-tetra-H4Kac) were utilised to detect histone acetylation in late trophozoite stage by immunofluorescence assay (IFA). Hoechst staining and PfH4Kac immunolabeling identified histone acetylation in late trophozoite nuclei **(Fig.5c)**. The acetylation pattern can be seen in fluorescence pictures, where SAHA increases and S1P decreases the intensity labelling of acetylated H4 (**Fig. 5d**). In cell analysis was done to analyze the binding of S1P in parasite. S1P agarose bead pull down assay was performed. First nuclear lysate was prepared according to protocol mentioned in material method section (**Fig. 5e**, To see if host S1P binds to parasite HDAC-1 in the cell, we first incubated nuclear lysate with S1P beads, then with HDAC-1 antibodies. S1P conjugated beads showed a prominent band in immunoblotting assays using recombinant HDAC-1, however non-conjugated beads as a negative control did not. **Fig. 5f**). S1P-containing beads were employed in a pull-down assay to confirm that S1P interacts with parasite HDAC-1 but not erythrocyte HDAC-1. *Pf*HDAC-1 antibody detects HDAC-1 in nuclear lysate but not erythrocytes. **(Fig.5g)**. Next coomassie dye-stained gel shows a band at ∼52 kDa band of HDAC-1 in lane 1 pulldown by S1P conjugated agarose bead and lane 2 shows input of nuclear lysate proteins used to pull down protein which interact with S1P in nucleus (**Fig. 5h**).

Lipid dot blot experiment revealed HDAC-1 and S1P binding. 50ng, 100ng, 200ng, and 400ng S1P were detected on nitrocellulose membrane. When incubated with parasite nuclear lysate, dark spots of varied sizes developed on S1P dots, confirming HDAC1-S1P binding. Absence of a spot in the phosphatidic acid (PA)-bound region showed HDAC1’s selectivity for S1P. (**Fig. 5i**). Overall, these data indicate that S1P interacts with HDAC-1 in parasite nuclei, where it modulates the activity of HDAC-1, as demonstrated by the preceding results.

**Figure 5:**
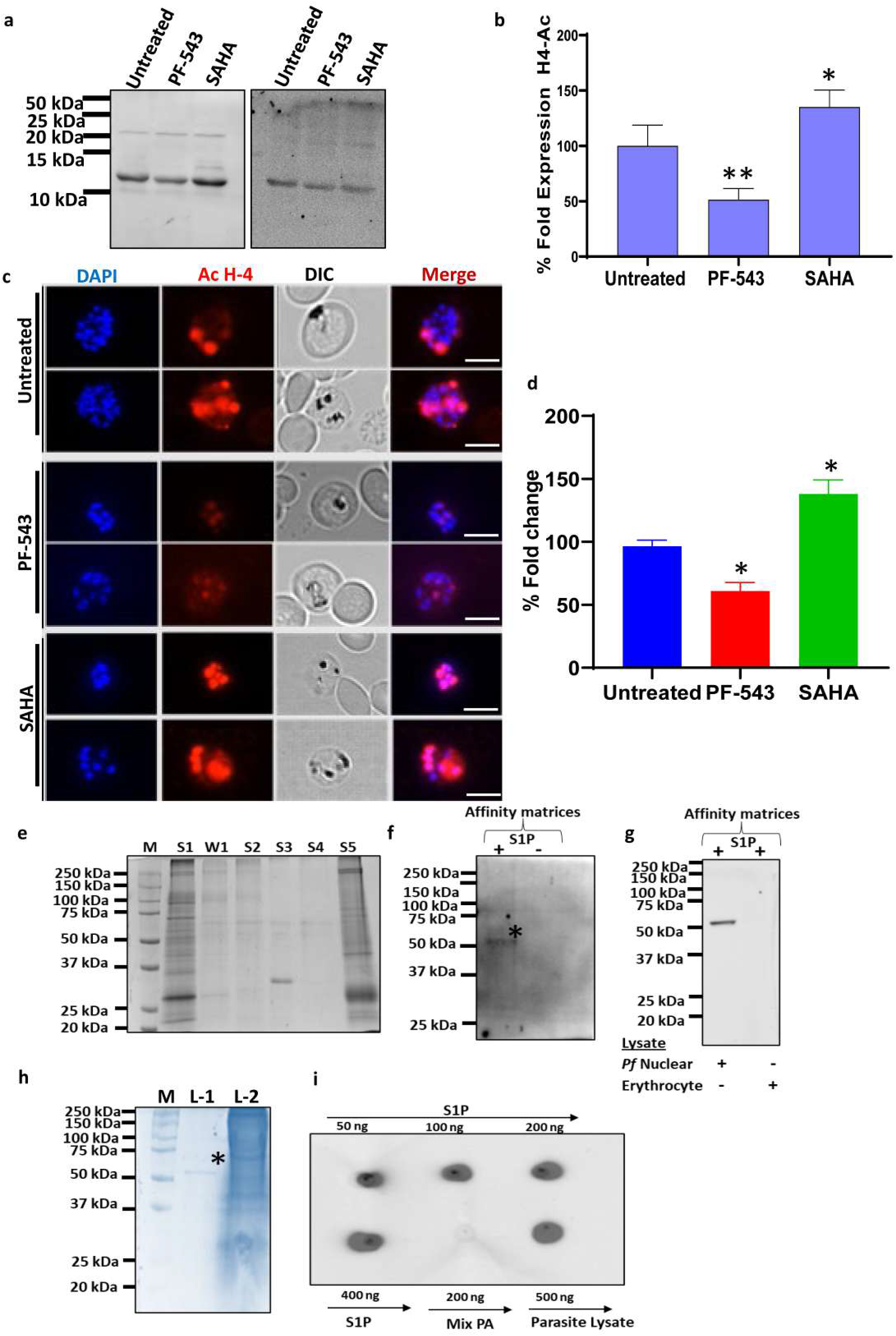
Host Sphk-1 inhibition by PF-543 regulate HDAC mediated acetylation of histone. **a-b)** Histone acetylation in nuclear extracts isolated from trophozoites (20– 28 hpi) treated with DMSO (0.005%), PF-543 (IC90) and SAHA was detected via immunoblotting using rabbit anti-H4Kac4 antibodies and mice anti-H4. Quantification of acetylation was done by measuring the band intensities via Image J for five different experiments; (n=5 independent experiments, *p*-value:< 0.0001), the values were normalized with the band intensities of anti-H4 used as loading control. **c-d)** Histone acetylation and hyper-acetylation following treatment with PF-543 and SAHA for 6 h at 37◦C in trophozoite stages were subjected to Immunofluorescence analysis via immunolabeling using rabbit anti-H4Kac4 antibodies (Red). Nuclei were stained by Hoechst nuclear stain (blue). Scale Bar, 5 μm. Fluorescence micrographs demonstrate altered histone acetylation level in parasite following treatment with PF-543. Bar graph denotes the differential mean fluorescence intensity (MFI) denoting H4Kac4level in parasite (n=3 independent experiment, *p*-value: < 0.0001) **e)** Cytoplasmic and nuclear fractions isolated from parasite were subjected to SDS-polyacrylamide gel electrophoresis. **f)** Pull-down of endogenous HDAC1 across asexual blood stage parasite using S1P agarose beads. Elutes obtained after washing S1P beads were subjected to SDS-polyacrylamide gel electrophoresis and the detection was done using immunoblot analysis. **g)** Presence of HDAC1 was detected only in parasite nuclear lysate. On a contrary, erythrocyte lysate showed no HDAC1 specific band. **h)** SDS-PAGE gel showing a band specific for rPfHDAC with an expected size of 52kDa in the elute fraction obtained from pull down assay (Lane 1). Lane 2 represents crude parasite lysate used for the experiment. **i)** Lipid dot blot with the indicated amounts of S1P, PA and crude parasite lysate were spotted onto Nitrocellulose Membrane filter. Binding of PfHDAC-1was detected with mice anti-PfHDAC1 polyclonal antibody. PfHDAC-1 showed significant binding with S1P. No binding was observed between PfHDAC and the negative control PA. PfHDAC-1 binding to different lipid concentrations was determined by immunoblot spot density. Values are given as arbitrary units (mean ± SEM; n = 3 independent experiments with proteins, \**p* ≤ 0.05).

### Downregulation of S1P by inhibition of host SphK-1 results in the deregulation of genes during parasite development

PfHDAC1 protein regulates several important biological pathways, including host cell entry/egress, cellular signaling, hemoglobin metabolism and the cell cycle. Given that these are processes associated with parasite development and the progression of infection in host cells^30^, we decided to observe the effects of host S1P lipid on PfHDAC-1 mediated gene regulation.

To determine if S1P binds PfHDACs and regulates gene expression in P. falciparum, we identified genes disrupted after asexual stage treatment with PF-543 and SAHA (positive control). Trophozoites were treated with PF-543 and SAHA for 3 h, and total RNA was extracted for comparison. At 3 h post treatment, total RNA was extracted from infected erythrocytes, and the resulting cDNA was subjected to real-time PCR, using α-tubulin, CPPUF, ETMP-4, PFMC-2TMI, PFMC-2TMII, GA27/25, PVMPS16, MDG1, PHISTB, CPPUF, CP47, AT2, ETMP10, PFEMPII, PFEMPI-II, RPL7, AIPAIP, H4,AP2 specific primers as reported in the **(Supplementary table 1).** RNA derived from untreated infected erythrocytes served as a negative control. The inhibition of host erythrocyte SphK1 by specific inhibitor PF-543, showed a profound decrease in the expression of HDAC regulated genes; α-tubulin (∼ 90%, p ≤ 0.01), CPPUF (∼ 76%, p ≤ 0.03), ETMP-4 (∼ 88%, p ≤ 0.02), PFMC-2TMI (∼ 47%, p ≤ 0.05), PFMC-2TMII (∼60%, p ≤ 0.02), GA27/25 (∼ 50%, p ≤ 0.05), PVMPS16 (∼ 75%, p ≤ 0.005), MDG1(∼ 80%, p ≤ 0.03), PHISTB (∼ 75%, p ≤ 0.04), CPPUF (1) (∼ 44%, p ≤ 0.16), AT2 (∼ 95%, p ≤ 0.04), ETMP10 (∼ 70%, p ≤ 0.01), PFEMPII (∼ 65%, p ≤ 0.17), PFEMPI-II (∼ 80%, p ≤ 0.02), RPL7 (∼ 80%, p ≤ 0.01), AIPAIP (∼ 95%, p ≤ 0.03), HH4 (∼ 57%, p ≤ 0.05)**(Fig6 a)**. The previous studies suggest alpha tubulin expression as a potential transcriptional marker of HDAC-1 inhibition in malaria parasites^30 20^. Importantly, 90% mRNA down regulation of alpha tubulin expression was observed **Fig 6a**).

**Figure 6:**
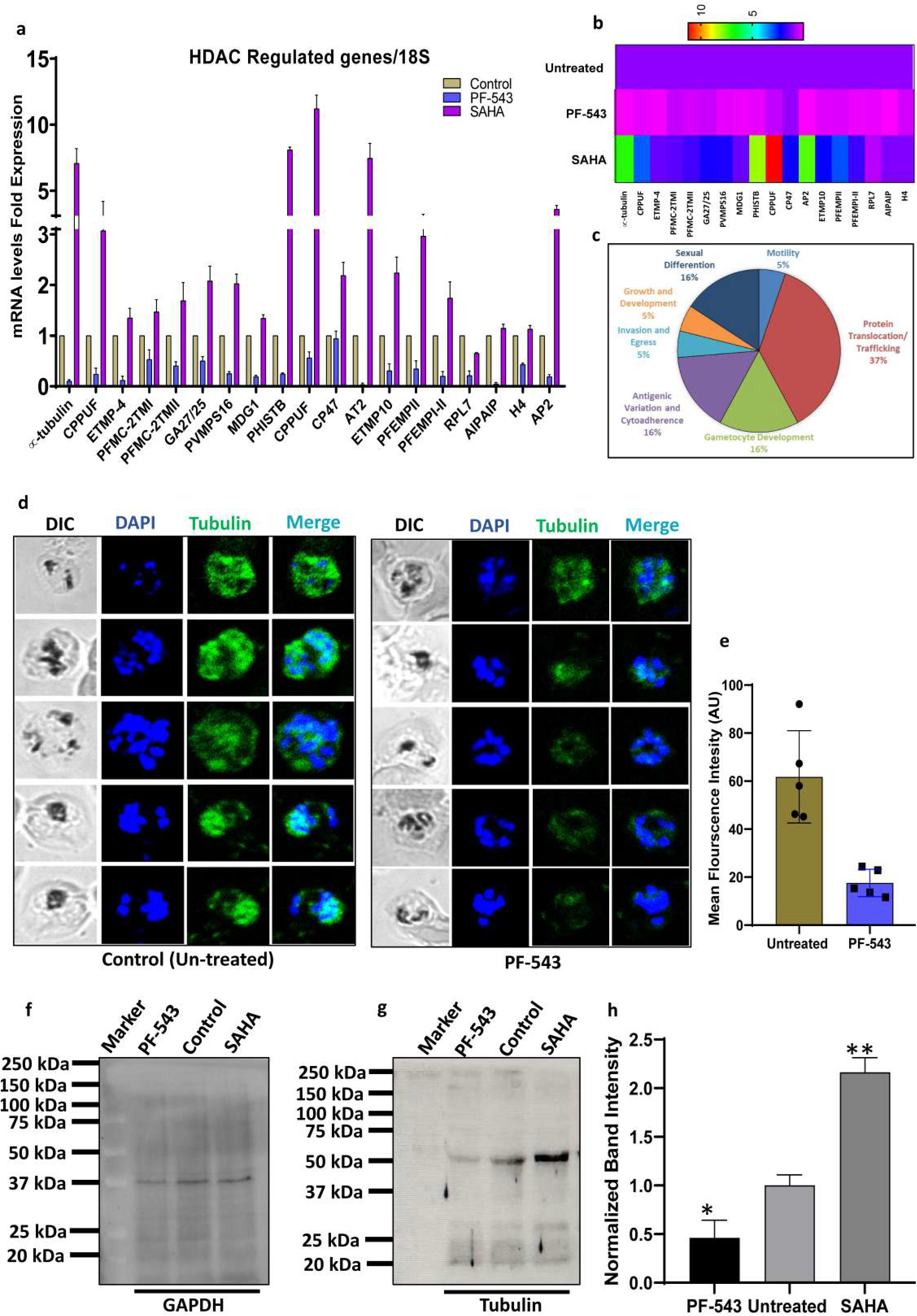
S1P mediated HDAC-1 regulation leads to change in global transcriptional response of *P. falciparum*. **a)** Shown is the Quantitative RT-PCR results carried out on 18 *P. falciparum* genes. From a total of 18 genes, 8 genes were found to be up-regulated in both biological replicates for SAHA and downregulated in PF-543 treated sample. Relative fold change in expression levels were calculated using 2−ΔΔCt method where 18S rRNA gene was used as internal control. **b).** The total number of genes with altered expression is shown graphically and through heat map. **c)** Genes commonly or uniquely regulated are shown in Pie chart. Data are the combined replicates for each compound to show genes commonly up- or down-regulated for each compound and treatment time. **d-e)** Immunofluorescence analysis exhibiting alteration of *P. falciparum* microtubule dynamics upon PF-543 mediated SphK-1 inhibition. Parasites at ring and early trophozoite stages were treated with PF-543 for 6 hr, permeabilized, stained with rabbit anti-α-tubulin antibody followed by Alexa Fluor 488 secondary antibody (red). Nuclear material was stained with DAPI (blue). Representative bar graph displayed changes in mean fluorescence intensities. (n=2 individual experiments, ∗p ≤ 0.05). **f-h)** Immunoblot analysis to further validate the differential expression of α-tubulin in parasites treated with PF543 (1 µM) and SAHA (2µM). The Protein was probed with mice polyclonal anti-His antibody and GAPDH was taken as loading control. Bar graph denotes the relative level of α-tubulin in PF-543 and SAHA treated parasites with respect to untreated control. (n=3 individual experiments, ∗p ≤ 0.05)

These results collectively suggest Sphingosine 1 Phosphate (S1P) as an endogenous regulator of PfHDAC. The heat map displays relative gene expression after PF-543 and SAHA treatment. It shows elevated and downregulated genes. (**Fig 6 b**). A pie chart classifies the PF-543 and SAHA-regulated genes by function. These genes are involved in protein trafficking, translocation, virulence, sexual commitment, and gametocyte formation. (**Fig 6 c**).

As tubulin is a highly specific indication of HDAC regulation, it can be used as a biomarker. Next, late trophozoites were incubated with 1μM PF-543 and 2.5 μM SAHA for 8 hours. After treatment, the cells were labeled with anti-tubulin antibody and analysed using fluorescence microscopy; the same sample was then processed for immunoblotting following saponin lysis. Following treatment with PF-543, the images demonstrated a drop in the fluorescence intensity of tubulin. (**Fig. 6 d**). The fluorescence intensities were plotted as bar graph by measuring ten individual cells using Image J for the treated and untreated cells indicating the same (**Fig. 6 e**). Immunoblot analysis of parasites treated with 1 μM PF-543 and 2.5 μM SAHA for 8 h showed a similar effect (**Fig. 6 f, g**). There was a considerable drop in global -tubulin following treatment with PF-543 and an increase after treatment with SAHA, which strongly validates the immunofluorescence data. The GAPDH gene was used as the positive control. The bar graph depicts the normalised alpha-tubulin-1 band intensity. (**Fig. 6h**).

### Inhibition of erythrocytic SphK-1 by PF-543 inhibit growth of malaria parasite and gametocyte in *in vitro*

Sphk-1 inhibitor P543 was evaluated for antimalarial effectiveness against asexual human malaria parasite in vitro to determine the link between host SphK1 activity and parasite growth. 3D7 ring-stage parasites were treated to a range of PF-543 concentrations for 72 h to determine IC_50_. Parasitaemia was assessed 72 h post-treatment using SYBR Green I fluorescence assay, and the average of three independent experiments was graphed. (**Fig.7a**). As seen in blood smears, PF-543 severely reduced parasite growth from rings to trophozoites, stopping the parasite at the ring stage. The appearance of ’pyknotic body’ after treatment halted parasite growth. (**Fig.7 b**). P543 showed substantial anti-malarial activity against 3D7, with an IC_50_ of 470 nm. These results support the concept that PF-543 impairs de-novo S1P production in host erythrocytes. Infected erythrocytes prevented the parasite from using the host’s S1P pool, which slowed growth. Next, the gametocidal activity of PF-543 was also evaluated in *P. falciparum* gametocytes. For this RKL-9 strain of *P. falciparum* was differentiated into sexual stages and treated with PF-543 at IC_50_ concentration derived from the asexual stages. Daily dose of PF-543 at early stage I cause the decline in gametocytaemia. Majority of the parasite formed pyknotic bodies by day 2. However, in DMSO treated control gametocyte develop till stage V as shown in (**Fig.7 c**). Result demonstrated that Sphk-1 inhibitor attenuate gametocyte development at early stages by modulating the S1P level **(Fig.7d).** After PF-543 therapy, one of the genes, AP-2, which is critical for gametocyte maturation, is downregulated. AP-2 has little influence on asexual development, but it’s important for gametocyte maturation, which could explain why PF-543 impairs gametocyte development.

**Figure 7:**
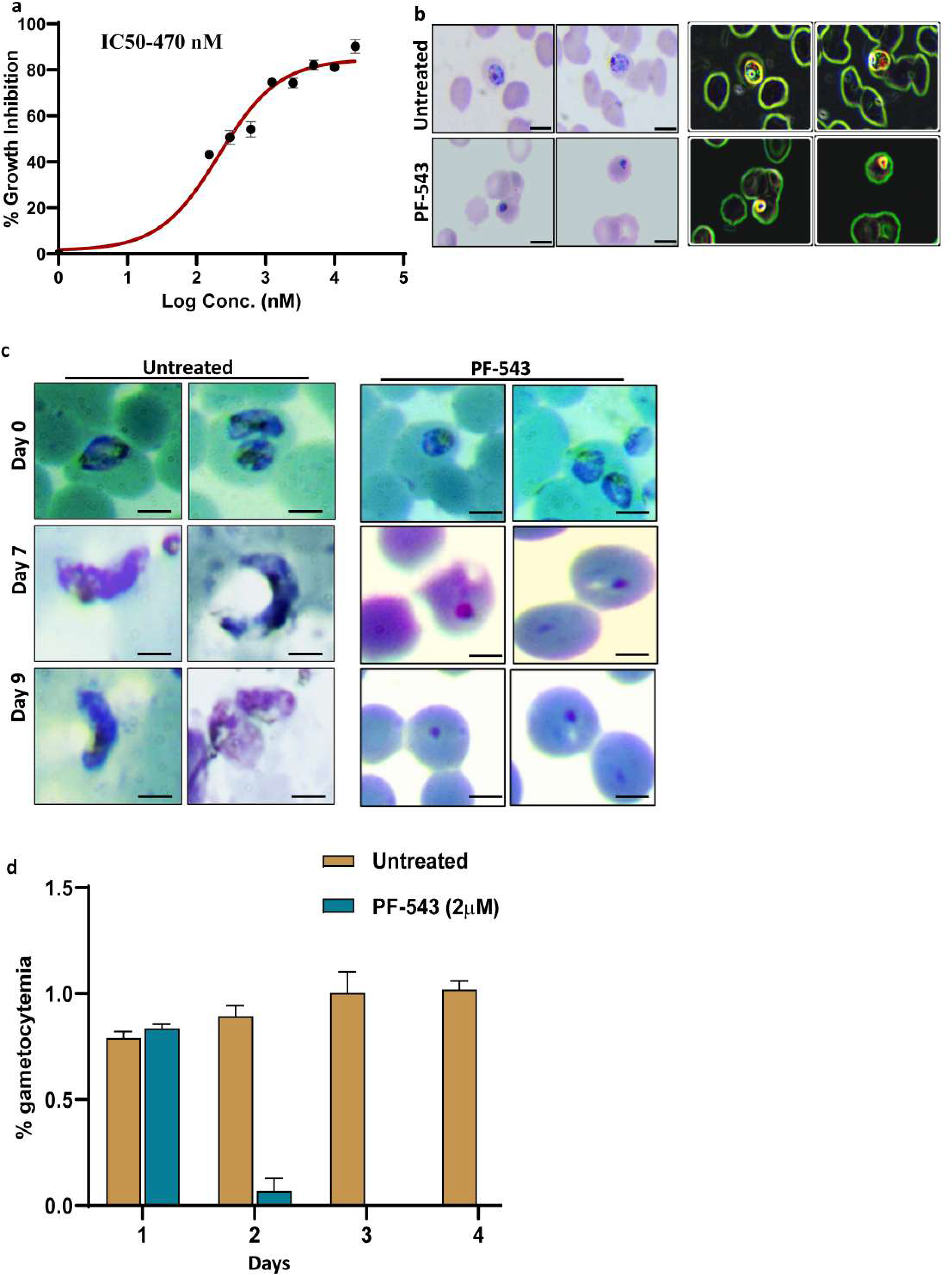
Inhibition of host SphK-1 blocks parasite growth and gametocytemia. **a)** Erythrocytes infected with *P. falciparum* strains 3D7, Dd2 and R539T were treated with different concentrations of PF-543 (0–8 µM) for 72 h and the percentage inhibition of parasite growth was evaluated, as represented in the graph. An IC50 value of 470 nM was determined for the parasite growth inhibition by PF-543. (n=3 independent expreiments, ∗p ≤ 0.05) **b)** Shown are the Giemsa stained smears of parasites treated with PF-543 with respect to the untreated control. **c)** Light microscope images of Giemsa stained parasites evincing inhibition of *P. falciparum* gametocyte progression following PF-543 treatment. Occurrence of pyknotic bodies demonstrates growth inhibition upon treatment. Scale bar-5μM**. d)** Bar graph representing effect of PF-543 on gametocyte formation. A significant decline in percent gametocytemia can be observed in PF-543 treated parasites as compared to no treatment control. (n=3 independent experiments. ∗p ≤ 0.05)

### Effects of SphK-1 inhibition by PF-543 on Early Sporogonic Stages

The lipid profiles of infected erythrocytes are characteristic for the parasite life cycle and maturity stages of gametocytes. The major membrane lipid class (phospholipids) decreased during gametocyte development. However, lipidomic analysis reveals an enrichment of sphingolipids and ceramides in gametocytes ^55, 56^. To assess the effect of low S1P level in host cell due to the inhibition of Sphk-1 by P543 on gametogenesis. PF-543 were tested for their activity against the gametocyte stage of *P.berghei* parasites. PF-543 was given intraperitoneally to phenyl hydrazine-treated mice to examine its effects on *P.berghei* gametocytes. In two-day *PF-543*-treated mice, gametocyte density was lowered by 50-60%. (**Fig 8a**. Representative, Giemsa-stained smear showed the healthy gametocyte morphology in DMSO treated mice. In contrast, immature and altered gametocyte morphology was observed PF-543 treated mice (**Fig.8 b)**. A significant decrease in gametocyte numbers was observed in treated mice evaluated in these studies.

**Figure 8:**
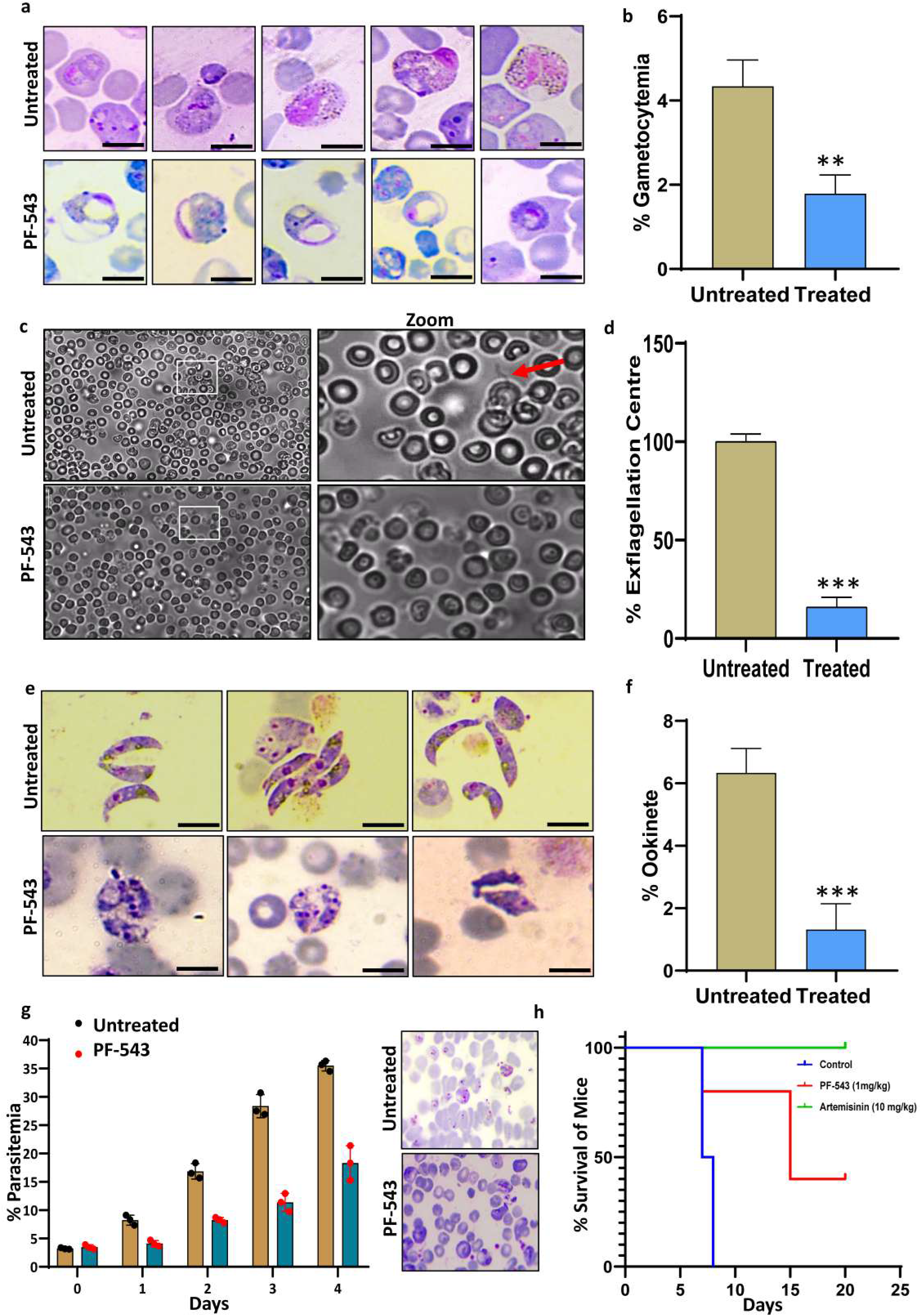
Effect of PF-543 treatment on sporogonic cycle and its effect on *P. berghei* infected mice survival. **a-b)** Giemsa stained smears of *P.berghei* showing gametocyte formation and morphology upon treatment with PF-543 as compared to untreated control. Bar graph showing a significant inhibition of gametocyte growth upon treatment with PF-543. (n=3 independent experiments, ∗p ≤ 0.05) **c-d)** Light microscopic images showing a reduction in exflagellation of *P.berghei* gametocytes following treatment with PF-543. Bar graphs representing a significant inhibition of gametocyte mediated exflagellation centre formation. (n=3 independent experiments, ∗p ≤ 0.05). **e-f)** Light microscope images of giemsa stained ookinetes showing growth inhibition upon PF-543 treatment. Bar graph representing the relative decrease in percent ookinete formation upon PF-543 treatment. (n= 3 independent experiments, ∗p ≤ 0.05)**. g)** Inhibitory effect of PF-543 on parasite growth confers enhanced survival in mice represented by a decline in percentage of parasite in blood. Untreated mice, on a contrary showed high parasitemia. **h)** Administration of PF-543 (1mg/kg) to mice resulted in their increased survival as compared to untreated control. Artemisinin (10mg/kg) was taken as positive control.

Next, We performed an ex vivo exflagellation assay with P. berghei infected RBCs taken from infected mice in order to evaluate the effect of PF-543 on male ex-flagellation. (**Fig. 8c**) These centres were counted at 40X magnification after DMSO and PF-543 were added. Male gametes are generated from infected erythrocytes after 15–20 minutes of ex-flagellation. To differentiate the mobile male gametes, light microscopy used. Comparing PF-543 to an untreated DMSO control, the amount of ex-flagellation centres was significantly reduced by PF-543. **(Fig.8d).** This data strongly implicated the role of S1P during exflagellation.

The in vitro conversion of P. berghei mouse malaria gametocytes to ookinetes was also observed after PF-543 treatment. This assay measures the effects of compound on gamete development, fertilisation, and ookinete formation to anticipate parasite transmission blocking throughout the insect’s life cycle. After 24 h post-treatment, Giemsa smears were made and observed under light microscope which demonstrated fully developed elongated morphology of ookinete in DMSO treated control group, while morphologically immature or partially developed ookinetes were observed in the treated group **(Fig.8e)**. Ookinete numbers were counted in each of the untreated and treated groups and represented as a bar graph (**Fig. 8f**). PF-543 reduced ookinete production by 80% compared to control, demonstrating S1P-mediated ookinete maturation. Epigenetic control may be important for stage-specific Plasmodium gene expression. S1P modulates HDAC’s activity. Altering S1P levels deregulated epigenetic control mediated by HDAC during the ookinete stage, which affected ookinete development after PF-543 treatment.

The results of in vitro antiplasmodial potency of PF-543 further encouraged us to evaluate their efficacy in vivo in mice infected with murine malaria parasite P. berghei. ANKA strain. The result showed that in treated mice parasitemia significantly reduced to 60% when compared with vehicle control (**Fig.8 9**). After three days of drug treatment mice were observed for further seven days to calculate the mean survival time and result showed that the treatment of compound increased the survival rate of P. berghei infected mice. Mice from the vehicle control group died within seven days of infection (**Fig.8 h**). While mice from the treated group survived the infection for more than 12 days. Also, the drug treatment was well tolerated by all mice as no significant side effects were observed. The result suggested that host SphK-1 inhibition can abolish intra-erythrocytic growth of *P. berghei* in in vivo. Epigenetic regulation has been suggested to play a major role in the stage specific gene expression during the Plasmodium life cycle. The model depicts the role of sphingosine 1 phosphate (S1P) as an endogenous regulator of *Pf*HDAC activity. Sphk-1 is responsible for the production of S1P inside the host cell. In the presence of its inhibitor PF-543 level of S1P goes down because of that S1P is no longer be able to bind *Pf*HDAC. In the absence of their interaction, HDAC cause the deacetylation of its downstream genes, resulting in downregulation of genes involved in regulation of plasmodium life cycle regulation, including asexual stage and sexual stage (**Fig. 9**).

**Figure 9:**
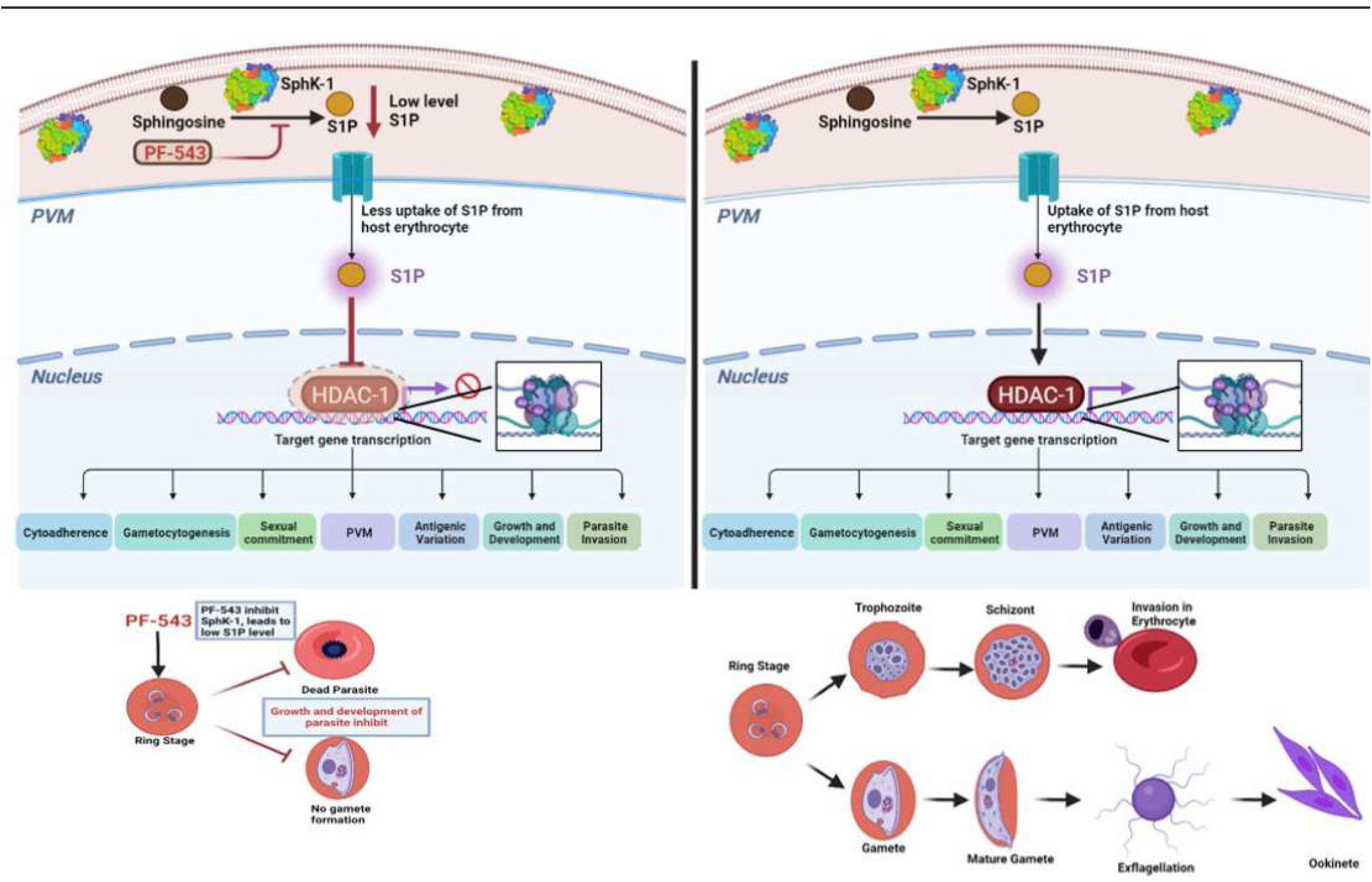
Schematic representation of S1P mediated epigenetic modulation of parasite survival pathways.

## Discussion

Our study found a previously unknown role for sphingolipid signaling in regulating the parasite critical gene HDAC-1, which controls the expression of several parasite genes. First, we show that S1P reprograms PfHDAC-1’s activity after engaging with it. This finding was unknown. We also show how S1P deregulates PfHDAC-1 activity, leading to an imbalance in the global transcriptome, and how this imbalance affects the parasite’s development. *Plasmodium falciparum* uses both its de novo pool and the host cell lipid repertoire during development and proliferation in ABS, signalinga metabolic connection that might be leveraged for developing antimalarials targeting the parasite’s metabolic co-dependency^50, 55, 57, 58^. Due to the lack of S1P degrading enzymes (S1P lyase and S1P phosphohydrolase) and its ability to acquire sphingosine from the environment, erythrocytes are the major reservoir for circulating S1P^59, 60^. S1P is a sphingolipid metabolite implicated in cell proliferation, survival, motility, angiogenesis, vascular maturation, immunity, and lymphocyte trafficking^61–63^. S1P interacts to histone deacetylases HDAC1 and HDAC2 in mammalian and suppresses their enzymatic activity, blocking the removal of acetyl groups from histone tail lysine residues^41^. HDACs are direct intracellular targets of nuclear S1P and regulate gene expression epigenetically. Parasites uptake S1P from the host pool because they lack Sphk1 isoform, but how this affects epigenetic regulation is unknown. Here, it has been demonstrated that S1P has an intracellular role in parasite epigenetic reprogramming. First, cloned pfHDAC-1 and purified protein for MST analysis of S1P binding **(Fig1)**. we molecular docked S1P to HDAC. S1P docks strongly with a binding energy comparable to that of SAHA, a potent HDAC inhibitor **(Fig2)**. We found that endogenous HDACs in parasite nuclear extracts bind to S1P immobilised on agarose beads. S1P binds selectively to His-tagged HDAC1 and endogenous HDAC from parasite nuclear lysate. HDAC and PA did not interact, demonstrating S1P binds solely to HDAC1 (**Fig. 2,5**). HDACs remove acetyl groups from histone lysine residues, compacting chromatin, and repressing transcription. Fluorescence-based HDAC activity assay and immunoblot analysis was used to determine if S1P binding affected HDAC activity. Our investigation confirmed that S1P binds to HDAC’s active site and suppresses its activity. S1P reduced HDAC1 activity by 50%, compared to SAHA’s 95%. Neither sphingosine nor LPA affected HDAC1 (**Fig.2, 5)**. NBD-sphingosine was utilized to detect S1P in nuclei. Infected erythrocytes incubated with NBD-Sph exhibited NBD-S1P in the parasite’s nucleus by spectrophotometry and fluorescence microscopy. NBD-S1P modulates HDAC-1 activity in the parasite’s nucleus, according to the study (**Fig.4**).

Lysine acetylation of histones is a critical component of an epigenetic indexing system demarcating transcriptionally active chromatin domains. Histone acetyltransferases (HATs) and HDACs maintain the delicate, dynamic equilibrium in acetylation levels of nucleosomal histones. Inhibiting HDACs activity will out-turn a dramatic increase in global acetylation of histones and subsequently a general induction of basal transcription. Our immunoblot and immunofluorescence data of H4 manifested the acetylation status of H4 in the presence of PF-543 and SAHA. As presumed, the acetylation of H4 was reduced in the presence of PF-543 and increased dramatically in SAHA as compared to the control (**Fig. 5**). In light with these observations, we performed quantitative RT-PCR assay to highlight the expression patterns of HDAC targeted genes in the presence of PF-543 and SAHA. Our data was found to correlate with the transcriptomics data reported in a recent publication by Kanyal et.al ^30^.

After HDAC-1 knockdown, genes involved in protein translocation and trafficking, gametocyte formation, sexual differentiation, and antigenic variation showed downregulation, correlating with our observations after lowering S1P. (**Fig 6**). This was also very much in agreement with recent study in HDAC KO cells^30^. However, we wished to establish the control of HDAC by S1P in cells via an experimental genetic technique; however, a recent article illuminates the crucial role of HDAC-1 in parasites. In addition, genetic manipulation cannot be utilized to illustrate the effect of S1P-mediated signaling utilizing knockout HDAC-1, which Kanyal et al. demonstrated is essential for parasite formation and growth ^30^. In order to confirm the effects of PF-543 and SAHA on the asexual blood stages of the parasite, we compared the -tubulin expression profile throughout these stages. Immunofluorescence and an Immunoblot assay indicated that PF-543-treated samples exhibited lower -tubulin expression than SAHA-treated samples relative to the untreated control. The level of S1P is associated with HDAC activity. (**Fig.6**). Furthermore, we have also shown that manipulating HDAC1 activity with potent Sphk-1 inhibitor PF-543, ensued reduction in parasite growth in 3D7 strain. By forming “pyknotic bodies,” this inhibitor concentration suppressed parasite proliferation (**Fig 7**).

Next, we tested Sphk-1’s effect on *P. falciparum* gametocytogenesis. PF-543 inhibited gametocyte progression from stage I to stage V in-vitro. sphk-1 inhibition affects gametocyte development (**Fig 7**). Ex-vivo investigation of PF-543 on *P. berghei* gametocytes in mice showed abnormal gametocytes. PF-543 inhibit gametocyte development by reducing S1P production. Low S1P levels can’t suppress HDAC activity, which downregulates a gametocyte-related gene. *Ex-vivo* exflagellation tests using *P.berghei* infected erythrocytes evaluated S1P’s role in male gamete exflagellation. PF-543-treated parasites had 80% less exflagellation centre than untreated parasites. This inhibitor is effective against both the asexual and sexual phases of the malaria parasite, reducing the possibility of transmission to mosquitoes and other community members. (**Fig 8**).

In-vivo investigation on *P.berghei* infected mice confirmed that 1mg/kg PF-543 reduced parasitemia, gametocytemia, and survival in mice. This work showed *Pf*HDAC1’s essential role in *P. falciparum* intraerythrocytic development. Our studies show that HDAC1 controls multiple parasite developmental processes. Thus, S1P binding to *Pf*HDAC1 suggests their interaction within the parasite nucleus to regulate critical functions. This study introduces SphK-1/S1P signaling in erythrocytes as an alternative mechanism for *P. falciparum* survival and differentiation. We failed to knocking down of host SphK-1 in BEL-A erythroid progenitor cell to make terminally differentiated red blood cells devoid of SPhK-1, but shows decrease in the late stage of terminal differentiation, hence production of erythrocyte in vitro is not possible which is consistent with the study led by Yong et al^64^. SPHK1 inhibition increases erythroid cell apoptosis in vitro, according to a recent study that demonstrated the anti-apoptotic function of S1P. To demonstrate the genetic effect of SphK-1/S1P signalingon parasite growth and development, knocking down SphK-1 in erythroid progenitor cells is therefore not an option. Therefore, the best way to demonstrate the role of S1P signaling in parasite epigenetic reprogramming was to utilize the SphK-1-specific inhibitor PF-543 that is specific to SphK-1 and in clinical trials^65–69^.

Prior to our investigation, the role of S1P in Infected erythrocytes remained a complete mystery. Our in vitro and in vivo research identified a previously unknown intracellular role for S1P in interacting with PFHDAC-1 and changing its epigenetic state. This finding presents a novel mechanism of lipid–protein interaction in the infected erythrocyte with *P. falciparum* for adaptation of the parasite, reveals a novel key regulatory function of intracellular S1P in parasite physiology, and reveals new therapeutic opportunities to improve target-based therapeutic approaches.

## Methods

### Cultivation of *P. falciparum-*infected erythrocytes and growth inhibition assay

*P. falciparum* 3D7 strain was cultured in RPMI 1640 (Invitrogen, Carlsbad, CA, USA) supplemented with 27.2 mg/L hypoxanthine (Sigma-Aldrich, St. Louis, MO, USA), 2 gm/L sodium bicarbonate (Sigma-Aldrich, St. Louis, MO, USA) and 0.5 gm/L AlbuMax I (Gibco, Grand Island, NY, USA) using O+ human erythrocytes, under mixed gas environment (5% O_2_, 5% CO_2_ and 90% N_2_) as described previously ^53^. For the assessment of half-maximal drug concentration for inhibition of malaria survival (IC_50_) values, synchronized *P. falciparum* infected erythrocyte cultures were used at late-ring or early trophozoite stage (18–24 h post infection (hpi)) at a parasitemia of 1%.

### Cloning of Histone deacetylase 1 (HDAC) for recombinant protein generation

Nucleotide sequence encoding full length sequence of *P. falciparum* HDAC (1-1380 bp) was selected for recombinant protein generation as a fusion protein with an N-terminal 6x-Histidine-tag using the vector pET-28 a(+) vector . DNA fragment for the gene cloning was PCR amplified from genomic DNA (Pf3D7) using the following primer pair: HDAC BamHI_FP: 5’-CGC<u>GGATCC</u>ATGTCTAATAGAAAAAAGGTTGC-3’,and HDAC_XhoI_RP: 5’-CCG<u>CTCGAG</u>TTAATATGGTACAATAGATTGATCC-3 with Phusion™ High-Fidelity DNA Polymerase (Thermo Scientific, US). The amplified DNA fragment was purified with QIAquick Gel Extraction Kit (Qiagen) using the manufacturer’s protocol. Purified HDAC insert and the expression vector pET-28 a(+) were digested with BamHI/XhoI restriction enzymes (New England Biolabs, UK), and were ligated over night at 16 °C using T4 DNA ligase (New England Biolabs, UK). The ligation mix was transformed into *E. coli* DH5-α competent cells and positive clones were screened by colony PCR followed by confirmation of the cloned plasmid with BamHI/XhoI restriction digestion and further transformed in BL21 DE3 *E. coli* expression strain for HDAC recombinant protein expression.

### Isolation of nuclear fraction

Nuclear isolation and extraction was carried out based on previous method ^54^, with minor modifications. Infected RBCs containing trophozoites and schizonts were lysed with saponin and washed three times with 1X PBS. The obtained parasite pellet was treated with a hypotonic cytoplasmic lysis buffer (20mM HEPES (pH 7.9), 10mM KCl, 1mM EDTA, 1mM EGTA, 0.65% NP-40, 1mM DTT, 1mM PMSF, 1X PIC) for 5 min on ice. The sample was centrifuged at 2000g for 5 min to collect the cytoplasmic extract (Fraction 1). The pellet was rinsed four times with hypotonic cytoplasmic lysis buffer and suspended in low salt buffer (20mM HEPES (pH 7.9), 0.1M KCl, 1mM EDTA, 1mM EGTA, 1mM DTT, 1Mm PMSF, 1X PIC) for 20min at 4ᵒC with constant agitation. The supernatant was saved after 3 minutes of centrifuging at 13,000 rpm (Fraction 2). Pellet washed twice in low salt buffer and chromatin dissolved in DNaseI-digestion buffer (20Mm Tris-HCl (pH 7.5), 15mM NaCl, 60mM KCl, 1mM CaCl_2_, 5mM MgCl_2_, 5mM MnCl_2_, 300mM sucrose, 0.4% NP-40, 1mM DTT, 1mM PMSF, 1X PIC) containing 100-500U DNaseI for 20min at RT under constant agitation. The sample was then centrifuged at 13,000rpm for 3min and the supernatant was collected (Fraction 3). The pellet was then washed twice in low salt buffer and solubilised in a high salt buffer (20mM HEPES (pH 7.9), 1M KCl, 1mM EDTA, 1mM EGTA, 1mM DTT, 1mM PMSF, 1X PIC) for 20min at 4ᵒC under constant agitation. The supernatant was further saved by centrifuging the suspension at 13,000rpm for 3min (Fraction 4). The pellet containing insoluble nuclear matrix fraction was washed twice in high salt buffer and finally solubilised in the SDS extraction buffer (2%SDS, 10mM Tris-HCl (pH 7.5)) for 20min at RT under constant agitation followed by centrifugation at 13,000rpm for 15min. The supernatant (Fraction 5) was collected and saved. All fractions were run on an SDS-PAGE gel and immunoblotted to evaluate their quality.

### Pull down

Approximately 100 ul of S1P-agarose beads, were washed twice with lysis buffer. The fraction 2 of nuclear lysates were incubated with beads overnight at 4ᵒC with constant gentle rotation. Protein-bound beads were washed by wash buffer (10 mmol HEPES pH 7.4, 150 mmol NaCl, 0.25% NP-40) for six times. Washed beads were added to 100ul of Laemmli buffer (Sigma-Aldrich) and heated at 100ᵒC for 5 min. Eluted proteins were separated by SDS–polyacrylamide gel electrophoresis and transblotted on nitrocellulose membrane. Anti-HDAC polyclonal antibody from mice confirmed HDAC on S1P beads. Secondary antibodies coupled with horseradish peroxidase were utilised for ECL. Similarly, recombinant HDAC protein was isolated and electrophoresed on SDS-polyacrylamide gel before western blotting.

### Dot Blot assay using nuclear fraction of parasite lysate

S1P was dissolved in methanol: chloroform and blotted on a nitrocellulose membrane. For negative control, a single dot of mixed Phosphatidic acid was also blotted. The membrane was then blocked-in blocking buffer (3% skim milk in TBS) for 30min at RT. Membrane was washed once using TBS and incubated with nuclear fraction 2 obtained after parasite lysis, overnight at 4ᵒC. Three washes, one with TBS and two with TBST (TBS with 0.05% Tween-20) were then given and the membrane was probed for 1 h with mice raised anti-HDAC polyclonal primary antibody at a dilution of 1:2000. After another three washes with TBS and TBST, the membrane was incubated with the secondary antibody: anti-mice HRP (1:8000) in TBST containing 1% skim milk, for 1 h at RT. Following three washes using TBS and TBST, detection of protein and lipid binding was performed using ECL method.

### Dot Blot of r*Pf*HDAC and rH4

Lipid dot blot assay was performed to detect the binding interaction between S1P and rPfHDAC/rH4. S1P was dissolved in methanol: chloroform solution and dots with varying concentration of S1P were blotted on two nitrocellulose membranes. The dots were air-dried followed by blocking of the membrane in blocking buffer (3% skim milk in TBS) for 30min at RT. Membranes were washed once using TBS and incubated with rPfHDAC1 and rH4 separately overnight at 4ᵒC. Three washes, one with TBS and two with TBST (TBS with 0.05% Tween-20) were then given and the membranes were probed for 1 h with mice raised anti-HDAC1 and anti-H4 polyclonal primary antibody at a dilution of 1:2000. After another three washes with TBS and TBST, the membrane was incubated with the secondary antibody: anti-mice HRP (1:8000) in TBST containing 1% skim milk, for 1 h at RT. Following three washes using TBS and TBST, detection of protein and lipid binding was performed using ECL method

### Microscale thermophoresis

Binding affinities of S1P with the recombinant PfHDAC and hHDAC was evaluated by MST analyses, using Monolith NT.115 instrument (NanoTemper Technologies, Munich, Germany). Additionallly, binding interaction between SAHA and recombinant PfHDAC/hHDAC was also measured as positive control. Sample was prepared by mixing 389nM r*Pf*HDAC1 in Phosphate buffered saline, pH 7.4 supplemented with 0.05% Tween-20 to prevent sample aggregation, followed by labelling with 30uM Cysteine reactive dye (Monolith Protein Labelling Kit Red-Maleimide 2^nd^ Generation, NanoTemper), and incubated in dark at RT for 30 minutes. The labelled r*Pf*HDAC1 along with the buffer was added to the equilibrated column, followed by collecting elution fractions. The elution fraction with fluorescence counts less than 1,000 was taken further for interaction analysis. Increasing concentrations of S1P and SAHA (1mM to 0.01nM respectively) were titrated against the constant concentration of the labelled r*Pf*HDAC1. Samples were pre-mixed and incubated for 10 minutes at RT in the dark, followed by loading of the samples into the standard treated capillaries (K002 Monolith NT.115). Data evaluation was performed with the Monolith software (Nano Temper, Munich, Germany).

### HDAC activity screening

The Histone Deacetylase Activity Assay Kit (Fluorometric; ab156064) was used to measure recombinant *Pf*HDAC1 and in parasite lysate. Corning black, 96-well plates were used and the following groups were run: no enzyme control, solvent control, and 20uM S1Pand 5uM SAHA was used as a positive control. The principle of the assay relies on the deacetylation of the fluoro-substrate peptide by HDACs. Fluoro-deacetylated peptides are produced in the process, which can then be hydrolyzed by lysyl endopeptidases into lysine and 7-amino-4-methylcoumarin (AMC). AMC leads to an increase in fluorescence intensity. The enzyme mixture was prepared by adding 100 ng of the diluted enzyme to 300 µl of HDAC buffer. From the enzyme mixture, 40 µl was taken and mixed with 10 µl of test compound (final to 20 µM) and SAHA (5 uM) or vehicle (control). The final mixture (50 µl) was added to each well which was then pre-incubated at 37 °C for 10 min. The HDAC reaction was started by adding 50 µl of HDAC substrate: Lys (Ac)-AMC. The plate was incubated at 37 °C for 45 min. Trypsin stop solution (50 µl) was added to the well, and the plate was further incubated at 37 °C for 15 min to stop the reaction and reading was taken using varioskan LUX Multimode Microplate Reader (Thermo fisher, Massachusetts, USA).

### Quantification of NBD-S1P with SphK-1 inhibitor using microplate reader

For the fluorometric SPHK assay, fluorescent substrate was used and the conversion of NBD-sphingosine (NBD-Sph) to NBD-S1P in infected erythrocyte after PF-543 treatment was quantified in nuclear extract. 100 µl of infected erythrocyte suspension (1 × 109 cells) with 6 to 8% parasitemia in iRPMI media and 100 µl of iRPMI containing 10 µM NBD-Sph were incubated at 37 °C for 60 min. After incubation, lipids were extracted from nuclear extract of saponin lysed parasites. Then, 260 µl of methanol and 400 µl of chloroform/methanol (1:1) were added to the samples and thoroughly mixed. Subsequently, 16 µl of 7 M NH4OH, 400 µl of chloroform, and 300 µl of 1.5 M KCl were added to the samples and mixed thoroughly. The lipids were separated by centrifugation for 5 min at 17,000×g and 100 μl of the upper (aqueous) phase transferred to a black 96-well plate. The fluorescence intensities of the upper aqueous phases containing NBD-S1P in treated and untreated sample were measured at 530 nm with excitation at 485 nm in 96-well plates using varioskan LUX Multimode Microplate Reader (Thermo fisher, Massachusetts, USA).

### Immunofluorescence assay for histone acetylation

For immunofluorescence assays, thin smears of schizont or mixed stage parasites treated or untreated with PF-543 and SAHA were made on glass slides, air dried and fixed with methanol (ice cold) for 30 min at − 20 °C. Smears were blocked with 3% (w/v) bovine serum albumin (BSA) in phosphate buffer saline (PBS) containing blocking buffer (pH 7.4) for 30 min at room temperature (RT). Slides were probed with anti-H4Kac4 (Invitrogen, Carlsbad, CA, USA, 1:1000) mice and in blocking buffer at RT for 1 h. After washing, slides were incubated with Alexa Fluor 594 conjugated goat anti-mouse IgG (Molecular Probes, USA, 1:500) at RT for 1 h. After washing, the slides were mounted in Prolong Gold antifade reagent (Invitrogen, Carlsbad, CA, USA), viewed on a Olympus fluorescence microscopy. Further, images were processed via NIS-Elements software.

### Immunodetection of altered histone modifications and induced protein expression

Parasite infected erythrocytes were treated in the presence or absence of PF-543 and SAHAat trophozoite stage. Cells collected at 4 hours posttreatment were lyzed in saponin (0.05%) and washed three times with PBS. The isolated parasites were resuspended directly in SDS sample buffer, incubated at 100^0^C for 10 min and centrifuged at16,000g for 10 min. The supernatant was analyzed by 15% SDS-PAGE and transferred onto the PVDF membranes (Millipore) followed by blocking with 5% skim milk blocking buffer for 1 h at 4 °C. After washing, blots were incubated for 1 h with anti-H4Kac4 (1:3000), anti-histone 4, anti-alpha tubulin (Invitrogen, Carlsbad, CA, USA, 1:1000) mouse and rabbit in blocking buffer. Later, the blots were washed and incubated for 1 h with appropriate secondary anti-mouse and anti-rabbit (1:10,000) antibodies conjugated to HRP. Immunoblotted proteins were visualized by using the Clarity Western ECL substrate (Bio-Rad).

### qRT-PCR for gene expression analysis

Total RNA from *P. falciparum* infected erythrocytes were isolated at appropriate time point using the TRIZOL reagent (Invitrogen, Grand Island, NY) and quantified using a Nanodrop ND-1000 spectrophotometer (Thermo Fischer, USA). cDNA was prepared from one micrograms of RNase-free DNase treated total RNA using first-strand cDNA Synthesis Kit (Thermo Fischer Scientific, USA), as per manufacturer’s instructions, using random hexamer primers. PCR reactions were carried in Applied Biosystems, Real-Time PCR System (ABI, CA, USA) using PowerUp SYBR Green PCR Master Mix (Thermo Fisher Scientific, USA). The detail of the primers (sequences and annealing temperatures) used is given in **Supplementary Table S1**. Thermal profile for the real-time PCR was amplification at 50°C for 2 min followed by 40 cycles at 95°C for 15 sec, 60°C for 30 sec and 72°C for 1 min. Melting curves were generated along with the mean C_T_ values and confirmed the generation of a specific PCR product. Amplification of 18S was used as internal control for normalization. The results were expressed as fold change of control (Untreated samples (18S)) using the 2 ^-ΔΔ*CT*^method. Each experiment was done in triplicates and repeated three times. Statistical significance was determined by Student’s t-test analysis (P<0.05).

### In vitro antimalarial activity of *PF-543* against human malaria parasite

Strains of P. falciparum, 3D7 (chloroquine-sensitive) were used for the chemosensitivity tests. The drugs were placed in 96-wells flat-bottom microplates in triplicate at different concentration. Sorbitol synchronized cultures with 0.8-1% parasitemia and 2% hematocrit were aliquoted into the plates and incubated for 72 h in a final volume of 100 µL/well. Chloroquine was used as a reference compound. Parasite growth was determined with SYBR Green I based fluorescence assay. Briefly, after 72 h of incubation culture was lysed by freeze-thaw. Followed by addition of 100 µL of lysis buffer (20 mM Tris/HCl, 5 mM EDTA, 0.16% (w/v) saponin, 1.6% (v/v) Triton X) containing 1× SYBR Green I ((Thermo Fisher Scientific, Waltham, Massachusetts, US)). Plates were incubated in the dark at room temperature (RT) for 3 -4 h. *P. falciparum* proliferation was assessed by measuring the fluorescence using a Varioskan™ LUX multimode microplate reader (Thermo Scientific™) with an excitation and emission of 485 nm & 530 nm respectively. IC50 values were determined via non-linear regression analysis using GraphPad prism 8.0 software. The results were expressed as the percent inhibition compared to the untreated controls.

### Animal handling

The investigations followed rules from JNU’s Institutional Animal Ethics Committee (IEAC) and the Committee for Control and Supervision of Animal Experiments (CPCSEA). IAEC-JNU’s animal ethics committee approved the regulations for the use of animals in research and in experimental research under strict adherence to ethical standards. In the experiment, mice were housed under standard conditions of food, temperature (25 ± 3 °C), relative humidity (55 ± 10%) and illumination (12 h light/dark cycles) obtained from the Central Laboratory Animal Resources, Jawaharlal Nehru University, Delhi.

### In-vivo antimalarial activity of *PF-543* against rodent malaria parasite in mice

To check the antimalarial activity of PF-543 in vivo. Chloroquine-sensitive *Plasmodium berghie* ANKA strain (*Pb*ANKA) was used for the induction of malaria in the experimental BALB/c mice. Mice infected with *Pb*ANKA were used as the donor source. The donor infected mice with a parasitemia of 20 –30% were sacrificed, and blood was collected by cardiac puncture into heparinized tubes. The blood was subsequently diluted with 0.9% normal saline solution and the infection was induced by injecting 0.2 ml of the diluted blood containing 1 × 10^8^ parasitized erythrocytes via IP injection (20). Parasitemia was monitored daily by microscopic examination of Giemsa-stained thin blood smears, and was calculated using the following formula, % parasitemia = Number of infected erythrocytes × 100/ Total number of erythrocytes.

To check the antimalarial activity infected mice were divided into 3 group, each group comprise of 3 mice. One day post infection by *P. berghei*, one groups of mice were given compound PF-543 respectively via I.P route for 3 consecutive days at a dose of 1 mg/kg of body weight. Two control group were used in parallel: one was treated with artemisinin (10 mg/kg) and the other was treated with vehicle (DMSO). Doses were given once in a day. To evaluate the parasitemia in infected mice upon drug administration smear was made by collecting blood from tail of each mice regularly, stained with Giemsa 10% (v/v) and examined under the bright field microscope. Parasitemia was evaluated and the percent inhibition of parasite growth was calculated in relation to the untreated group.

### Effect of Sphk-1 inhibitor *PF-543* on *P. berghei* exflagellation ex-vivo

Mice were given phenylhydrazine 30 mg/kg intraperitoneally to evaluate PF-543 effect on P. berghei male gametocyte ex-flagellation. Four days post-treatment, mice were infected with 1.0×10^7^ *P.berghei* ANKA parasites through I.P. 3 days after infection, gametocytemia peaks. Five days later, 120 μL of infected blood was obtained from mice and mixed with complete RPMI. Infected RBCs were then treated with 2 μM of *PF-543*, for control infected RBCs were left untreated. Samples were kept at 37°C and incubated for 1 h. After incubation drug treated blood samples were washed and mixed immediately with 200 μL of ex-flagellation medium (RPMI1640 containing 25 mM HEPES, 20% FBS, 10 mM sodium bicarbonate and 50 mM xanthurenic acid at pH 8.0) and kept at 20°C for 15 min. Ex-flagellation centres were counted in 10-12 fields at 40X magnification.

### Effect of Sphk-1 inhibitor *PF-543* on *P. berghei* ookinete development ex-vivo

To evaluate the effect of these compound on ookinete maturation same proportional of *P.berghei* infected RBCs were obtained from phenyl hydrazine treated mice. Infected RBCs were then mixed with ookinete media (RPMI1640 containing 25 mM HEPES, 20% FBS, 10 mM sodium bicarbonate (Sigma-Aldrich) and 50 mM xanthurenic acid (Sigma-Aldrich) at pH 8.4), maintaining 10% hematocrit along with *PF-543* at 2 μM concentration for 21-24 h at 21°C. Ookinete development was followed by Giemsa-stained smears.

### Gametocyte maturation in microwell plates

To analyse gametocyte development in 96-well plates, asexual parasite cultures were synchronized by purifying schizonts percoll gradient-centrifugation and allowed to reinvade erythrocytes. After two rounds of reinvasion the cultures were treated for 72 h with NAG in order to clear residual asexual parasites and obtain a virtually pure gametocyte culture. Aliquots of 100 μL of synchronized gametocyte culture (typically at 2%–4% gametocytaemia), diluted to 1% HCT, were seeded in 96-well flat-bottomed plates in presence of *PF-543* at IC50 and 2x IC50 concentration respectively. For control gametocyte were left untreated. Media with appropriate concentration of *PF-543* was exchanged daily for continuous 12 days. Giemsa-stained blood smears were observed to see the effect of *PF-543* on gametocyte development.

## Acknowledgements

Funding from National Bioscience Award from the Department of Biotechnology (DBT), Government of India (Sanction No.BT/HRD/NWBA/39/04/2018 -19) is acknowledged. RKS and GK are recipients of fellowship from Council of Scientific and Industrial Research (CSIR), Government of India. SA acknowledges University Grant Commission (UGC) for doctoral research fellowship. EM recipients of SRA fellowship from Council of Scientific and Industrial Research (CSIR). MS acknowledges University Grant Commission (UGC) for doctoral research fellowship. The funders had no role in study design, data collection and analysis, decision to publish, or preparation of the manuscript.

## Author contributions

SS conceived the idea, designed the experiments, interpreted the results, wrote and edited the manuscript. RKS, SA and GK carried out the experiments, interpreted the results, analysed the experimental data, and wrote the manuscript. EM performed the qRT-PCR experiment and analysis. MS carried out cloning. RSH and AG assisted in mice experiment and microscopy.

