## Supplementary File for "Sphingosine-1-phosphate regulates Plasmodium histone deacetylase activity and exhibits epigenetic control over cell death and differentiation"

### SUPPLEMENTARY FILE 1

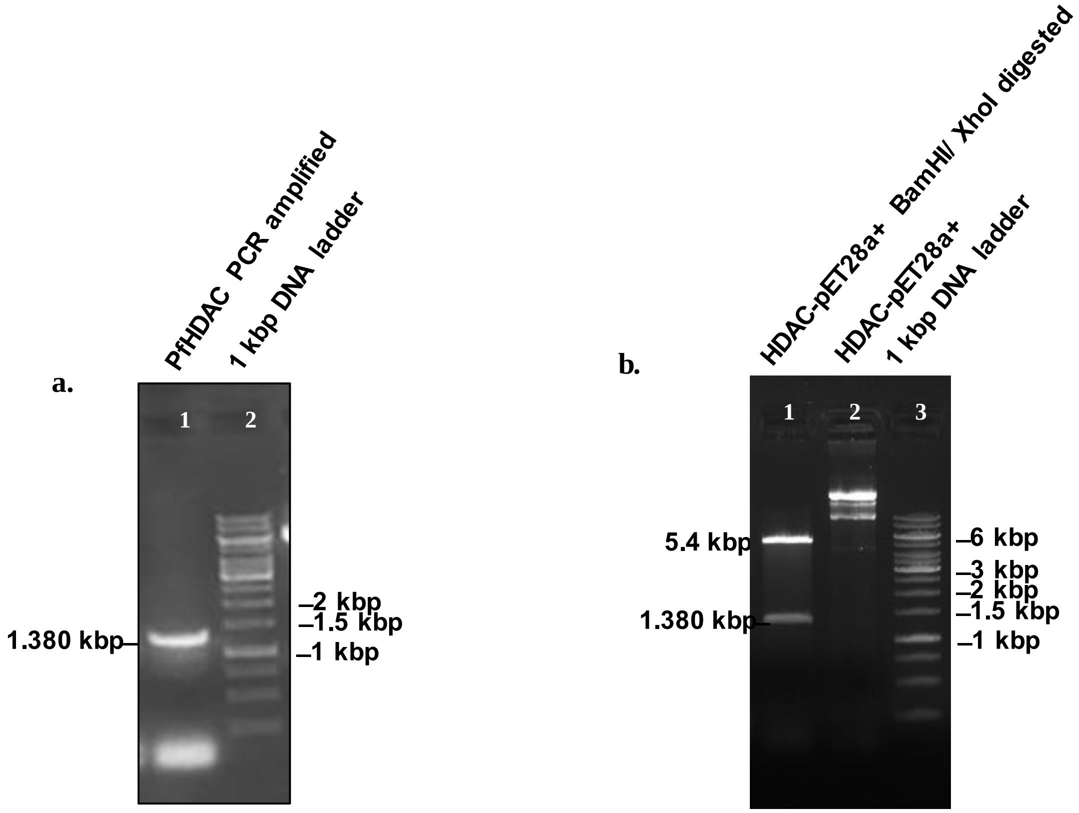


**Supplementary Fig 1. Cloning of *Plasmodium falciparum* Histone deacetylase-1 in pET28a+ vector. a.** Histone deacetylase-1 from *Plasmodium falciparum* was PCR amplified using Phusion™ High-Fidelity DNA Polymerase. Lane 1 represents the band specific for PfHDAC-1 gene at an expected size of 1380 bp. Lane 2 repesents 1kb DNA ladder*. **b.** Restriction digestion of the cloned pET28a+ plasmid vector with BamHI/XhoI restriction enzymes giving the fallout of HDAC-1 insert at 1380 bp. Lane 1 represents the digested plasmid obtained at a size of 5.4 kbp and the fallout of HDAC-1 insert at 1380bp. Lane 2 represents cloned pET28a+ plasmid vector with HDAC-1 insert. Lane 3 represents 1kb DNA ladder*. (*1kb DNA Ladder-SM#0331, thermo Scientific GeneRuler DNA Ladder mix.)

#### SUPPLEMENTARY FILE 2

| S.No. | Target genes | Forward Primer | Reverse Primer | Annealing Temp.(°C) |
|---|---|---|---|---|
| 1. | α-tubulin | 5′-TGAACATGGAATTCAACCGG-3′ | 5′- CGTCAACGACGGTGGGTTC-3′ | 61 |
| 2. | CPPUF | 5’TTCTTCCTCTCTTGAGCATTCC3’ | 5’TTTCATGTTGCTTTGATTTACG3’ | 61 |
| 3. | ETMP4 | 5’TCTTAGGTAGTGCTTTAGGTTTTGG3’ | 5’TGCTTTCATCTTTTTCGTCACC3’ | 61 |
| 4. | PFMC-2TMI | 5’TTTGGATTGTTTCAACGACCT3’ | 5’CATGTCAGGAAAATAACGAGCA3’ | 61 |
| 5. | PFMC-2TMII | 5’ATTTGGATTGTTTCAACGACCT3’ | 5’CATGTCAGGAAAATAACGAGCA3’ | 61 |
| 6. | GA27/25 | 5’GCCCTTGGATAAATTTGGAA3’ | 5’GGATCCTTGCTAAGGGTCATC3’ | 61 |
| 7. | PVMPS16 | 5’TTCTTCGCTTTTGCAAACCT3’ | 5’AAAGGCATTTTGTCAGCAGAA3’ | 61 |
| 8. | MDG1 | 5’TAGGAGCAAAAGCAGGTGAT3’ | 5’CCGTTTCTTCATTAGCATTTCC3’ | 61 |
| 9. | PHISTB | 5’TGAAGATACGCACTTGGATGA3’ | 5’TCTTTCAATTTTCCCACATCG3’ | 61 |
| 10. | CPPUF(1) | 5’GAAGCAAAAACGTGGACCAT3’ | 5’TGACGTCTTTCCCTTTTGCT3’ | 61 |
| 11. | 6-CP47 | 5’TTTAATCCCCTGACTAATGTTAAGC3’ | 5’ACTTCTTCGTATATTTCAGATGACC 3’ | 61 |
| 12. | AT2 | 5’ATGCAGTAGGTGGAGGTACAGG3’ | 5’CTGTCGATACTTGAGGAGATGG3’ | 61 |
| 13. | ETMP10.3 | 5’ATGAAGGTTTCTAGGCATACCG3’ | 5’AACGCTCTCTTATCATCATTTGC3’ | 61 |
| 14. | PFEMP1I | 5’TAGCCTGTGATGATTTTGAACC3’ | 5’GACGTTTCTTTTGTTGTTGTTCC3’ | 61 |
| 15. | PFEMP1II | 5’GTCCCCACATATTTCGACTACG3’ | 5’ACAAAAATCTTCTGCCCATTCC3’ | 61 |
| 16. | 60SRPL7-3 | 5’AGGTTCCCCCTTCAATTAACC3’ | 5’TTTTATCGGTTTTGGATTCAGG3’ | 61 |
| 17. | AIPAIP | 5’GAAAACGAGTTAAAAGAAAATGACG3’ | 5’CTGATGTATGGGATGAATAGCC3’ | 61 |
| 18. | HH4 | 5’GGTAAGGGAGGTAAAGGTTTGG3’ | 5’CTTCTTGCTAAACGTCTGATGG3’ | 61 |
| 19. | 18S | 5’- CCGCCCGTCGCTCCTACCG-3’ | 5’-CCTTGTTACGACTTCTCCTTCC-3’ | 55-60 |

**Supplementary Table 1**. Table showing the primer sequences of the gene panel which were studied for their regulation by *Pf*HDAC1 and SAHA through quantitative RT-PCR. The primer sequence of 18s rRNA gene which was used as positive control has also been mentioned in the table.

### SUPPLEMENTARY FILE 3

a.

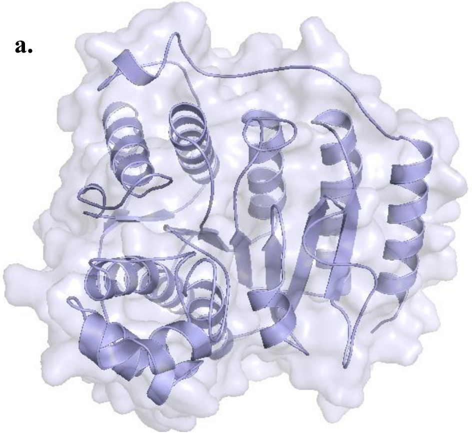

b.

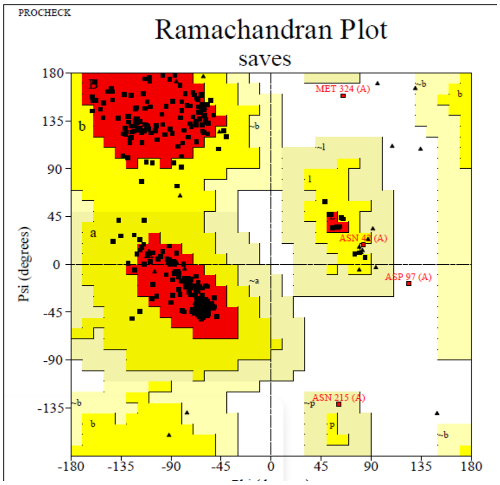


**Supplementary Fig 3. Molecular Docking analysis to study protein-protein interaction. a.** Energy minimized 3D protein model of *Pf*HDAC-1 used in the docking studies. **b.** Ramachandran plot of modeled structure of *Pf*HDAC-1 depicting 90.5% amino acid residues in the core region, 8.3% amino acid residues in the allowed region, 0.6% residues in the general region and only 0.6% residues in the disallowed region.

| | | | | | | | |
|---|---|---|---|---|---|---|---|
| **Hydrophobic Interactions** | | | | | | | |
| **Index** | **Residue** | **AA** | **Distance** | **Ligand Atom** | **Protein Atom** | | |
| 1 | 19A | TYR | 3.66 | 3003 | 137 | | |
| 2 | 19A | TYR | 3.36 | 3006 | 135 | | |
| 3 | 22A | ALA | 3.27 | 3002 | 160 | | |
| 4 | 77A | LEU | 3.6 | 3008 | 629 | | |
| 5 | 94A | GLU | 3.42 | 3013 | 782 | | |
| 6 | 99A | PRO | 3.69 | 3013 | 816 | | |
| 7 | 101A | PHE | 3.14 | 3010 | 830 | | |
| **Hydrogen Bonds** | | | | | | | |
| **Index** | **Residue** | **AA** | **Distance H-A** | **Distance D-A** | **Donor Angle** | **Donor Atom** | **Acceptor Atom** |
| 1 | 22A | ALA | 2.09 | 2.93 | 143.57 | 3024 [O3] | 159 [O2] |
| 2 | 22A | ALA | 1.92 | 2.81 | 152.07 | 3022 [O3] | 159 [O2] |
| 3 | 94A | GLU | 1.68 | 2.61 | 168.62 | 784 [O3] | 3015 [N3] |
| 4 | 94A | GLU | 1.83 | 2.61 | 130.16 | 3015 [N3] | 784 [O3] |

**Supplementary Table 2.** Table representing the amino acid residues involved in establishing the interaction between S1P and PfHDAC-1.

| Hydrophobic Interactions | | | | | | | |
|---|---|---|---|---|---|---|---|
| Index | Residue | AA | Distance | Ligand Atom | Protein Atom | | |
| 1 | 185B | GLU | 3.15 | 3684 | 1797 | | |
| 2 | 188B | TYR | 3.35 | 3684 | 1824 | | |
| 3 | 188B | TYR | 3.33 | 3687 | 1827 | | |
| **Hydrogen Bonds** | | | | | | | |
| **Index** | **Residue** | **AA** | **Distance H-A** | **Distance D-A** | **Donor Angle** | **Donor Atom** | **Acceptor Atom** |
| 1 | 144B | LYS | 3.09 | 4.03 | 153.58 | 1394 [Nam] | 3659 [O3] |
| 2 | 145B | SER | 3.06 | 3.87 | 136.84 | 1407 [Nam] | 3671 [O3] |
| 3 | 185B | GLU | 1.9 | 2.82 | 166.49 | 1800 [O3] | 3659 [O3] |
| 4 | 185B | GLU | 2.54 | 3.08 | 116.71 | 3671 [O3] | 1800 [O3] |
| 5 | 185B | GLU | 1.62 | 2.59 | 153.94 | 3662 [N3] | 1800 [O3] |

**Supplementary Table 3.** Table representing the critical amino acid residues involved in establishing the interaction between S1P and hHDAC-1.

| Hydrophobic Interactions | | | | | | | |
|---|---|---|---|---|---|---|---|
| Index | Residue | AA | Distance | Ligand Atom | Protein Atom | | |
| 1 | 155A | PHE | 3.88 | 3702 | 1509 | | |
| 2 | 209A | TYR | 3.73 | 3706 | 2048 | | |
| 3 | 209A | TYR | 3.35 | 3705 | 2050 | | |
| 4 | 210A | PHE | 3.35 | 3706 | 2059 | | |
| 5 | 210A | PHE | 3.74 | 3700 | 2060 | | |
| 6 | 210A | PHE | 3.88 | 3701 | 2061 | | |
| 7 | 276A | LEU | 3.03 | 3702 | 2670 | | |
| **Hydrogen Bonds** | | | | | | | |
| **Index** | **Residue** | **AA** | **Distance H-A** | **Distance D-A** | **Donor Angle** | **Donor Atom** | **Acceptor Atom** |
| 1 | 100A | ASN | 2.22 | 3.2 | 159.99 | 1014 [Nam] | 3691 [O2] |
| 2 | 104A | ASP | 2.59 | 3.42 | 137.02 | 3685 [N3] | 1047 [O2] |
| 3 | 104A | ASP | 2.26 | 3.09 | 144.2 | 3682 [O3] | 1048 [O3] |
| 4 | 104A | ASP | 2.3 | 2.8 | 112.82 | 1048 [O3] | 3694 [O3] |
| 5 | 104A | ASP | 1.93 | 2.8 | 151.92 | 3694 [O3] | 1048 [O3] |
| 6 | 153A | SER | 2.12 | 2.91 | 140.82 | 3692 [O3] | 1491 [O3] |
| 7 | 154A | GLY | 1.94 | 2.84 | 145.97 | 1493 [Nam] | 3694 [O3] |

**Supplementary Table 4.** Table representing the critical amino acid residues involved in establishing the interaction between S1P and hHDAC-2.

| Hydrophobic Interactions | | | | | | | |
|---|---|---|---|---|---|---|---|
| Index | Residue | AA | Distance | Ligand Atom | Protein Atom | | |
| 1 | 148A | PHE | 3.14 | 3671 | 1446 | | |
| 2 | 148A | PHE | 3.65 | 3672 | 1445 | | |
| 3 | 202A | TYR | 3.09 | 3666 | 1980 | | |
| 4 | 203A | PHE | 3.92 | 3659 | 1992 | | |
| 5 | 203A | PHE | 3.04 | 3665 | 1989 | | |
| 6 | 203A | PHE | 3.36 | 3670 | 1993 | | |
| 7 | 269A | LEU | 3.58 | 3669 | 2592 | | |
| 8 | 269A | LEU | 3.13 | 3668 | 2590 | | |
| 9 | 301A | TYR | 3.77 | 3673 | 2896 | | |
| **Hydrogen Bonds** | | | | | | | |
| Index | Residue | AA | Distance H-A | Distance D-A | Donor Angle | Donor Atom | Acceptor Atom |
| 1 | 138A | HIS | 2.32 | 3.33 | 171.92 | 1355 [Npl] | 3676 [O2] |
| 2 | 139A | HIS | 2.83 | 3.65 | 137.2 | 1368 [Npl] | 3676 [O2] |
| 3 | 147A | GLY | 2.33 | 3.3 | 159.22 | 3677 [Nam] | 1434 [O2] |
| 4 | 174A | ASP | 2.64 | 3.55 | 163.09 | 3679 [O3] | 1702 [O2] |
| 5 | 176A | HIS | 2.04 | 2.91 | 141.56 | 1719 [Npl] | 3676 [O2] |
| 6 | 299A | GLY | 3.62 | 4.02 | 105.22 | 2877 [Nam] | 3676 [O2] |
| 7 | 301A | TYR | 2.72 | 3.67 | 176.89 | 2899 [O3] | 3676 [O2] |
| **π-Stacking** | | | | | | | |
| Index | Residue | AA | Distance | Angle | Offset | Stacking Type | Ligand Atoms |
| 1 | 176A | HIS | 4.55 | 63.27 | 0.85 | T | 3664, 3665, 3666, 3667, 3668, 3669 |

**Supplemetary Table 4 :-** Table representing the critical amino acid residues involved in establishing the interaction between SAHA and *Pf*HDAC-1.

### SUPPLEMENTARY FILE 4

**a.**

#### XP_001352127.1 histone deacetylase 1 [Plasmodium falciparum 3D7]

Sequence ID: **Query_24109** Length: **449** Number of Matches: **1**

**Range 1: 3 to 398** Graphics ▼ Next Match ▲ Previous Match

| Score | Expect | Method | Identities | Positives | Gaps |
|---|---|---|---|---|---|
| 535 bits(1379) | 0.0 | Compositional matrix adjust. | 238/396(60%) | 312/396(78%) | 4/396(1%) |

```
Query  7    TRRKVCYYYDGDVGNYYYGQGHPMKPHRIRMTHNLLLNYGLYRKMEIYRPHKANAEEMTK  66
             R+KV Y++D D+G+YYYG GHPMKP RIRMTH+L+++Y LY+ ME+YRPHK++  E+T
Sbjct  3    NRKKVAYFHDPDIGSYYYGAGHPMKPQRIRMTHSLIVSYNLYKYMEVYRPHKSDVNELTL  62

Query  67   YHSDDYIKFLRSIRPDNMSEYSKQMQRFNVGE--DCPVFDGLFEFCQLSTGGSVASAVKL  124
            +H  +YI FL SI  +N  E++ Q++RFNVGE  DCPVFDGLF+F Q   G S+  A KL
Sbjct  63   FHDYEYIDFLSSISLENYREFTYQLKRFNVGEATDCPVFDGLFQFQQSCAGASIDGASKL  122

Query  125  NKQQTDIAVNWAGGLHHAKKSEASGFCYVNDIVLAILELLKYHQRVLYIDIDIHHGDGVE  184
            N    DI VNW+GGLHHAK SEASGFCY+NDIVL ILELLKYH RV+YIDID+HHGDGVE
Sbjct  123  NHHCADICVNWSGGLHHAKMSEASGFCYINDIVLGILELLKYHARVMYIDIDVHHGDGVE  182

Query  185  EAFYTTDRVMTVSFHKYGEYFPGTGDLRDIGAGKGKYYAWNYPLRDGIDDESYEAIFKPV  244
            EAFY T RVMTVSFHK+G+YFPGTGD+ D+G   GKYY+WN PL DG+ D+++  +FK V
Sbjct  183  EAFYVTHRVMTVSFHKFGDYFPGTGDITDVGVNHGKYYSWNVPLNDGMTDDAFVDLFKVV  242

Query  245  MSKVMEMFQPSAVVLQCGSDSLSGDRLGCFNLTIKGHAKCVEFVKSFNLPMLMLGGGGYT  304
            + K ++ ++P A+++QCG+DSL+GDRLG FNLTIKGHA+CVE V+S+N+P+L+LGGGGYT
Sbjct  243  IDKCVQTYRPGAIIIQCGADSLTGDRLGRFNLTIKGHARCVEHVRSYNIPLLVLGGGGYT  302

Query  305  IRNVARCWTYETAVALDT--EIPNELPYNDYFEYFGPDFKLHISPSNMTNQNTNEYLEKI  362
            IRNV+RCW YET V L+    E+P+++  NDY++Y+ PDF+LH+ PSN+ N N+ E+L +I
Sbjct  303  IRNVSRCWAYETGVVLNKHHEMPDQISLNDYYDYYAPDFQLHLQPSNIPNYNSPEHLSRI  362

Query  363  KQRLFENLRMLPHAPGVQMQAIPEDAIPEESGDEDE  398
            K ++ ENLR + HAPGVQ   +P D     +  DE +
Sbjct  363  KMKIAENLRHIEHAPGVQFSYVPPDFFNSDIDDESD  398
```

**b.**

#### XP_001347363.1 histone deacetylase 2 [Plasmodium falciparum 3D7]

Sequence ID: **Query_37627** Length: **2379** Number of Matches: **2**

**Range 1: 1167 to 1279** Graphics ▼ Next Match ▲ Previous Match

| Score | Expect | Method | Identities | Positives | Gaps |
|---|---|---|---|---|---|
| 55.8 bits(133) | 1e-11 | Compositional matrix adjust. | 30/113(27%) | 57/113(50%) | 5/113(4%) |

```
Query  204   YFPGTGDLRDIGAGKGKYYAWNYPLRDGIDDESYEAIFKPVMSKVMEMFQPSAVVLQCGS  263
             ++P TG   ++G  +G  + +N PL   G ++      +FK ++   ++E F+P  + + CG
Sbjct  1167  FYPRTGAKNELGEKEGYKFNINVPLEKGYNNCDVYYVFKYLLLPILEKFRPEFIFISCGF  1226

Query  264   DSLSGDRLGCFNLTIKGHAKCVEFVKSF-----NLPMLMLGGGGYTIRNVARC  311
             D+    D LG  NLT   +       +K F     N  ++++   GGY +   + +C
Sbjct  1227  DASINDPLGKCNLTHNLYQWMTFQLKHFANIFCNGRIILVLEGGYNLNYLPKC  1279
```

**Range 2: 956 to 1033** Graphics ▼ Next Match ▲ Previous Match ▲ First Match

| Score | Expect | Method | Identities | Positives | Gaps |
|---|---|---|---|---|---|
| 52.4 bits(124) | 1e-10 | Compositional matrix adjust. | 27/78(35%) | 41/78(52%) | 5/78(6%) |

```
Query  129   TDIAVNWAGGL---HHAKKSEASGFCYVNDIVLAILELLKYH--QRVLYIDIDIHHGDGV  183
             TDI  +A      HH  +S  SGFC  N+I +A   + K +  ++V   D D+HH +G
Sbjct  956   TDINCGFAAIRPPGHHCSRSHPSGFCIFNNISVACKYIFKKYGIRKVFIFDWDVHHDNGT  1015

Query  184   EEAFYTTDRVMTVSFHKY  201
             +E FY    V+  S H++
Sbjct  1016  QEIFYGDKDVLCFSIHRF  1033
```

c.

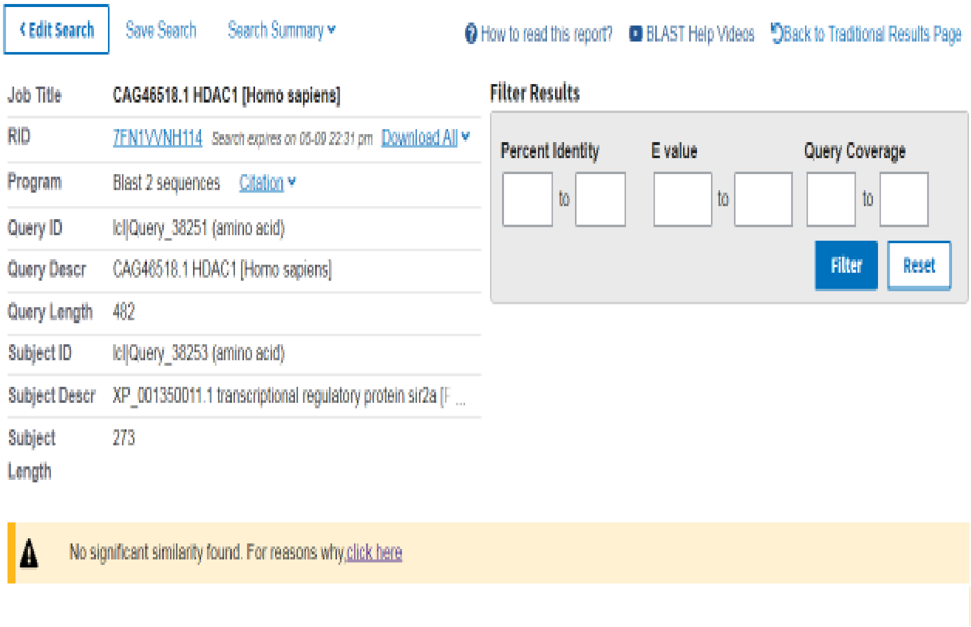


d.

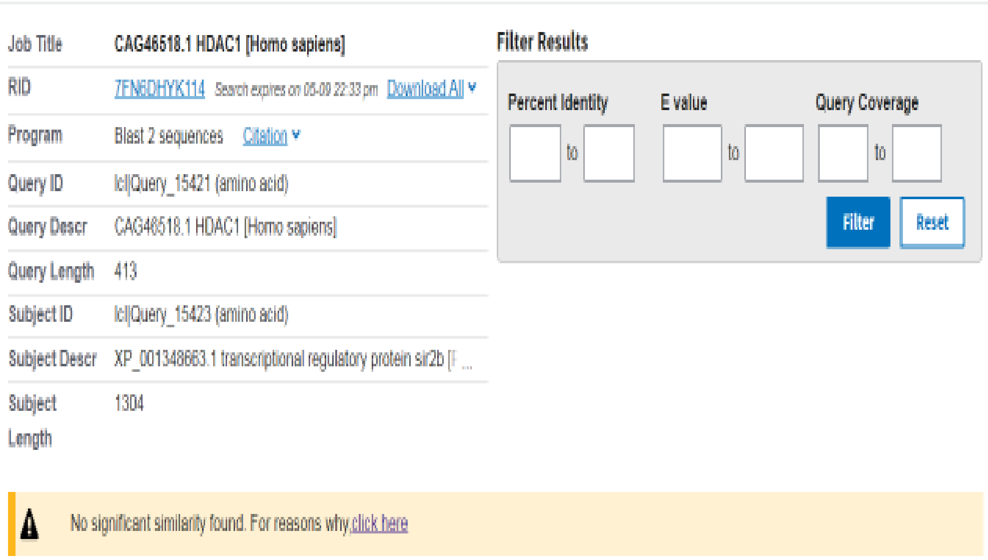


**Supplementary Fig 4. Sequence alignment data of human HDAC-1 with *Plasmodium falciparum* Histone deacetylase isoforms. a**. sequence alignment between hHDAC1 and *Pf*HDAC1 showing 60% sequence identity. **b.** Sequence alignment between hHDAC1 and PfHDAC2 showing 27% sequence identity. **c-d.** Sequence alignment of hHDAC1 with PfSir2a and PfSir2b showing no significant similarity.

**a.**

##### XP_001352127.1 histone deacetylase 1 [Plasmodium falciparum 3D7]

Sequence ID: **Query_62009** Length: **449** Number of Matches: **1**

Range 1: 4 to 396 Graphics ▼ Next Match ▲ Previous Match

| Score | Expect | Method | Identities | Positives | Gaps |
|---|---|---|---|---|---|
| 542 bits(1396) | 0.0 | Compositional matrix adjust. | 241/393(61%) | 313/393(79%) | 4/393(1%) |

```
Query  9    KKKVCYYYDGDIGNYYYGQGHPMKPHRIRMTHNLLLNYGLYRKMEIYRPHKATAEEMTKY  68
            +KKV Y++D DIG+YYYG GHPMKP RIRMTH+L+++Y LY+ ME+YRPHK+   E+T +
Sbjct  4    RKKVAYFHDPDIGSYYYGAGHPMKPQRIRMTHSLIVSYNLYKYMEVYRPHKSDVNELTLF  63

Query  69   HSDEYIKFLRSIRPDNMSEYSKQMQRFNVGE--DCPVFDGLFEFCQLSTGGSVAGAVKLN  126
            H  EYI FL SI  +N  E++ Q++RFNVGE  DCPVFDGLF+F Q   G S+ GA KLN
Sbjct  64   HDYEYIDFLSSISLENYREFTYQLKRFNVGEATDCPVFDGLFQFQQSCAGASIDGASKLN  123

Query  127  RQQTDMAVNMAGGLHHAKKSEASGFCYVNDIVLAILELLKYHQRVLYIDIDIHHGDGVEE  186
                D+ VNM+GGLHHAK SEASGFCY+NDIVL ILELLKYH RV+YIDID+HHGDGVEE
Sbjct  124  HHCADICVNMSGGLHHAKMSEASGFCYINDIVLGILELLKYHARVMYIDIDVHHGDGVEE  183

Query  187  AFYTTDRVMTVSFHKYGEYFPGTGDLRDIGAGKGKYYAVNFPMRDGIDDESYGQIFKPII  246
            AFY T RVMTVSFHK+G+YFPGTGD+ D+G   GKYY+VN P+ DG+ D+++  +FK +I
Sbjct  184  AFYVTHRVMTVSFHKFGDYFPGTGDITDVGVNHGKYYSVNVPLNDGMTDDAFVDLFKVVI  243

Query  247  SKVMEMYQPSAVVLQCGADSLSGDRLGCFNLTVKGHAKCVEVVKTFNLPLLMLGGGGYTI  306
             K ++ Y+P A+++QCGADSL+GDRLG FNLT+KGHA+CVE V+++N+PLL+LGGGGYTI
Sbjct  244  DKCVQTYRPGAIIIQCGADSLTGDRLGRFNLTIKGHARCVEHVRSYNIPLLVLGGGGYTI  303

Query  307  RNVARCNTYETAVALD--CEIPNELPYNDYFEYFGPDFKLHISPSNMTNQNTPEYMEKIK  364
            RNV+RCN YET V L+    E+P+++  NDY++Y+ PDF+LH+ PSN+ N N+PE++ +IK
Sbjct  304  RNVSRCNAYETGVVLNKHHEMPDQISLNDYYDYYAPDFQLHLQPSNIPNYNSPEHLSRIK  363

Query  365  QRLFENLRMLPHAPGVQMQAIPEDAVHEDSGDE  397
             ++ ENLR + HAPGVQ   +P D  + D  DE
Sbjct  364  MKIAENLRHIEHAPGVQFSYVPPDFFNSDIDDE  396
```

**b.**

##### XP_001347363.1 histone deacetylase 2 [Plasmodium falciparum 3D7]

Sequence ID: **Query_56541** Length: **2379** Number of Matches: **2**

Range 1: 1167 to 1279 Graphics ▼ Next Match ▲ Previous Match

| Score | Expect | Method | Identities | Positives | Gaps |
|---|---|---|---|---|---|
| 55.1 bits(131) | 2e-11 | Compositional matrix adjust. | 28/113(25%) | 57/113(50%) | 5/113(4%) |

```
Query  205   YFPGTGDLRDIGAGKGKYYAVNFPMRDGIDDESYGQIFKPIISKVMEMYQPSAVVLQCGA  264
             ++P TG   ++G  +G  + +N P+  G ++      +FK ++  ++E ++P  + + CG
Sbjct  1167  FYPRTGAKNELGEKEGYKFNINVPLEKGYNNCDVYYVFKYLLLPILEKFRPEFIFISCGF  1226

Query  265   DSLSGDRLGCFNLTVKGHAKCVEVVKTF-----NLPLLMLGGGGYTIRNVARC  312
             D+   D LG  NLT   +       +K F     N  ++++  GGY +  + +C
Sbjct  1227  DASINDPLGKCNLTHNLYQWMTFQLKHFANIFCNGRIILVLEGGYNLNYLPKC  1279
```

Range 2: 970 to 1033 Graphics ▼ Next Match ▲ Previous Match ▲ First Match

| Score | Expect | Method | Identities | Positives | Gaps |
|---|---|---|---|---|---|
| 51.2 bits(121) | 2e-10 | Compositional matrix adjust. | 23/64(36%) | 36/64(56%) | 2/64(3%) |

```
Query  141   HHAKKSEASGFCYVNDIVLAILELLKYH--QRVLYIDIDIHHGDGVEEAFYTTDRVMTVS  198
             HH  +S  SGFC  N+I +A   + K +  ++V   D D+HH +G +E FY     V+  S
Sbjct  970   HHCSRSHPSGFCIFNNISVACKYIFKKYGIRKVFIFDWDVHHDNGTQEIFYGDKDVLCFS  1029

Query  199   FHKY  202
              H++
Sbjct  1030  IHRF  1033
```

c.

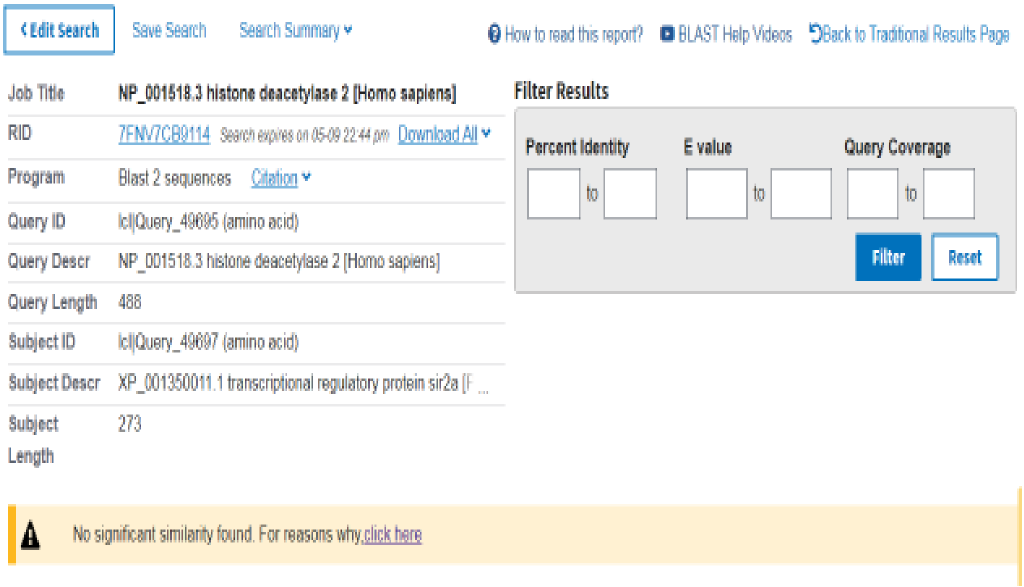


d.

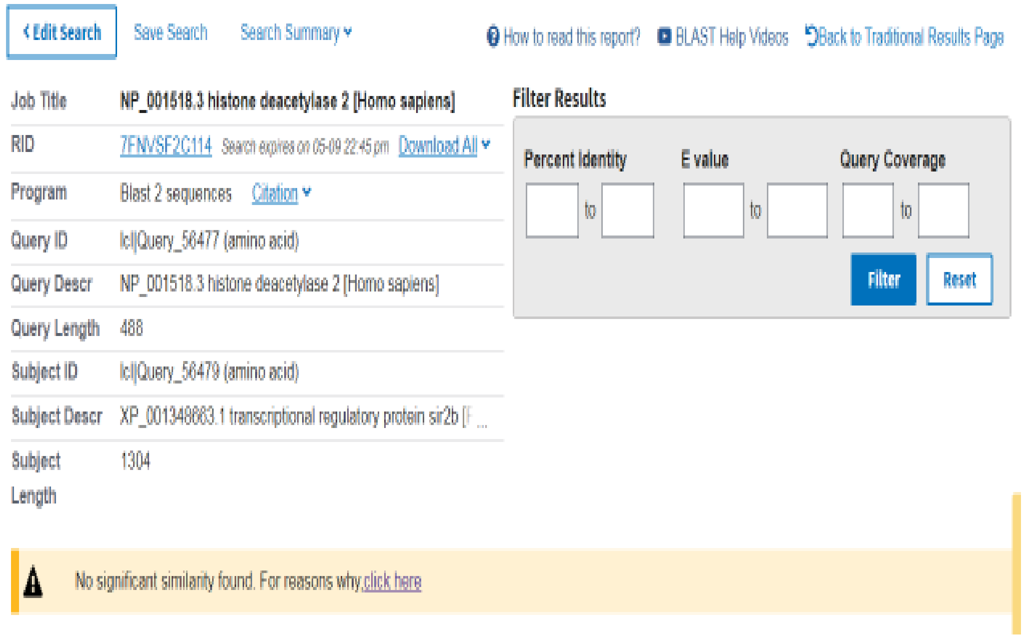


**Supplementary Fig 5. Sequence alignment data of human HDAC-2 with *Plasmodium falciparum* Histone deacetylase isoforms. a**. sequence alignment between hHDAC2 and *Pf*HDAC1 showing 61% sequence identity. **b.** Sequence alignment between hHDAC2 and PfHDAC2 showing 25% sequence identity. **c-d.** Sequence alignment of hHDAC2 with PfSir2a and PfSir2b showing no significant similarity.

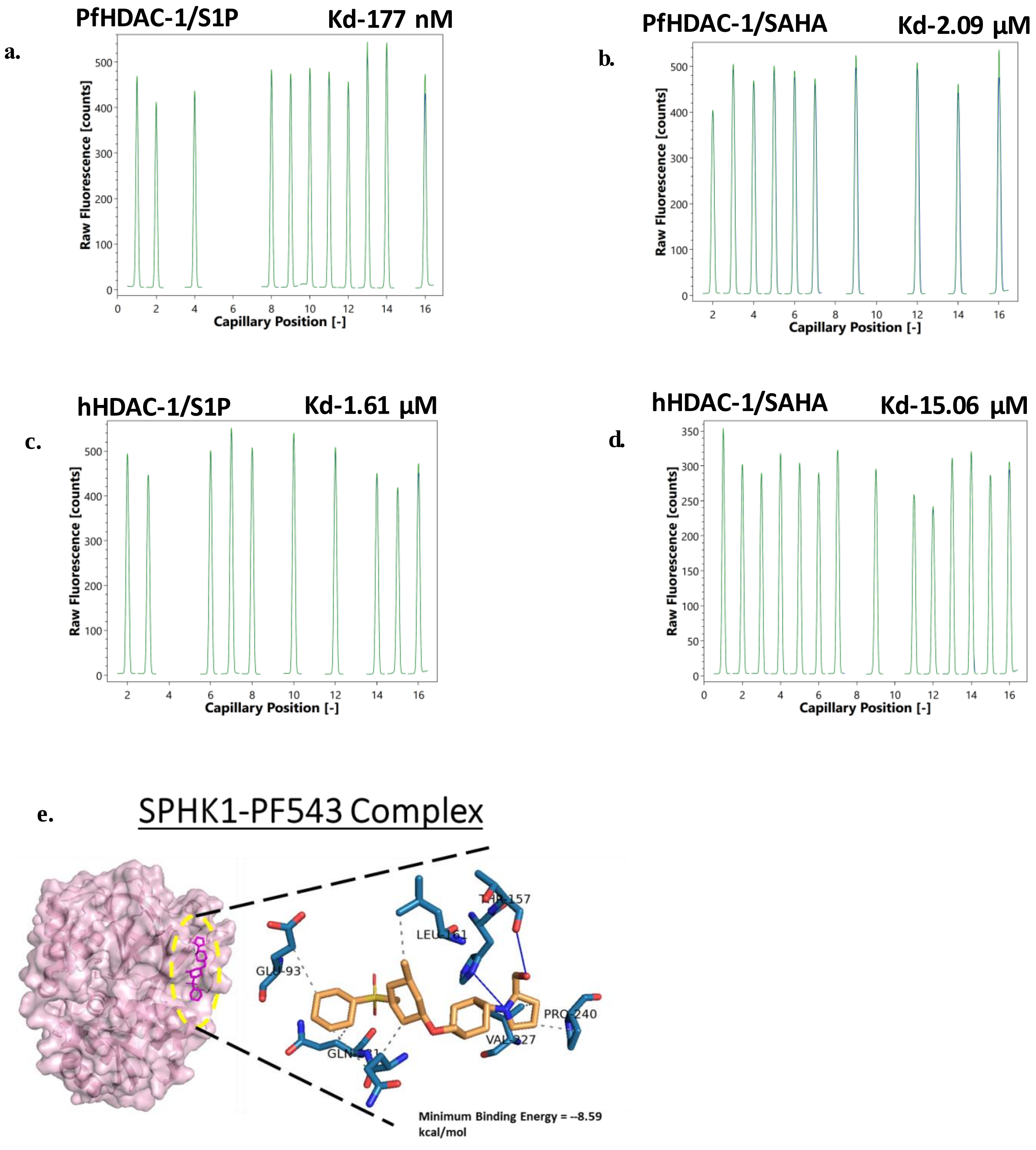


**Supplementary Fig 6.** Capillary scan profile of MST analysis of labelled **a.** PfHDAC-1 with S1P **b.** PfHDAC-1 with SAHA **c.** hHDAC-1 with S1P and **d.** hHDAC-1 with SAHA. **e.** Molecular docking analysis of hSphK-1 with its potent inhibitor PF-543.
